# TET dioxygenases localize at splicing speckles and promote RNA splicing

**DOI:** 10.1101/2025.03.07.641893

**Authors:** Florian D. Hastert, Jasmin Weber, Christina Bauer, Andreas Zhadan, Deepanshu N.D. Singh, Thomas C. Dix, Roland Arnold, Sergey Bessonov, Matthias Soller, Heinrich Leonhardt, M. Cristina Cardoso, Maria Arroyo

## Abstract

The dynamic regulation of RNA metabolism plays a crucial part in cellular function, with emerging evidence suggesting an important role for RNA modifications in this process. This study explores the relationship between RNA splicing and the TET dioxygenase activity, shedding light on the role of hm5C (RNA 5-hydroxymethylcytosine), and TET proteins, in RNA metabolism. Integrating data from mass spectrometry, AlphaFold structural modeling, microscopic analysis, and different functional assays including in vitro splicing, TET proteins were found to regulate splicing. We show that TET1, TET2, and TET3 interact with the splicing factors U2AF1 and U2AF2. Interestingly, TET dioxygenases localize in splicing speckles in mammalian and Drosophila cells. TET speckles association is RNA dependent, as it is TET interaction with splicing factors. Furthermore, in vitro splicing assays revealed that all three TET proteins promote splicing efficiency, and the oxidation of m5C to hm5C can restore splicing efficiency in vitro. The latter highlights the regulatory role of cytosine modifications in RNA metabolism. These findings provide insights into the complex interplay between RNA modifications and splicing, suggesting a multifaceted role for TET proteins in RNA metabolism beyond its canonical DNA demethylation function.

**Graphical abstract:** 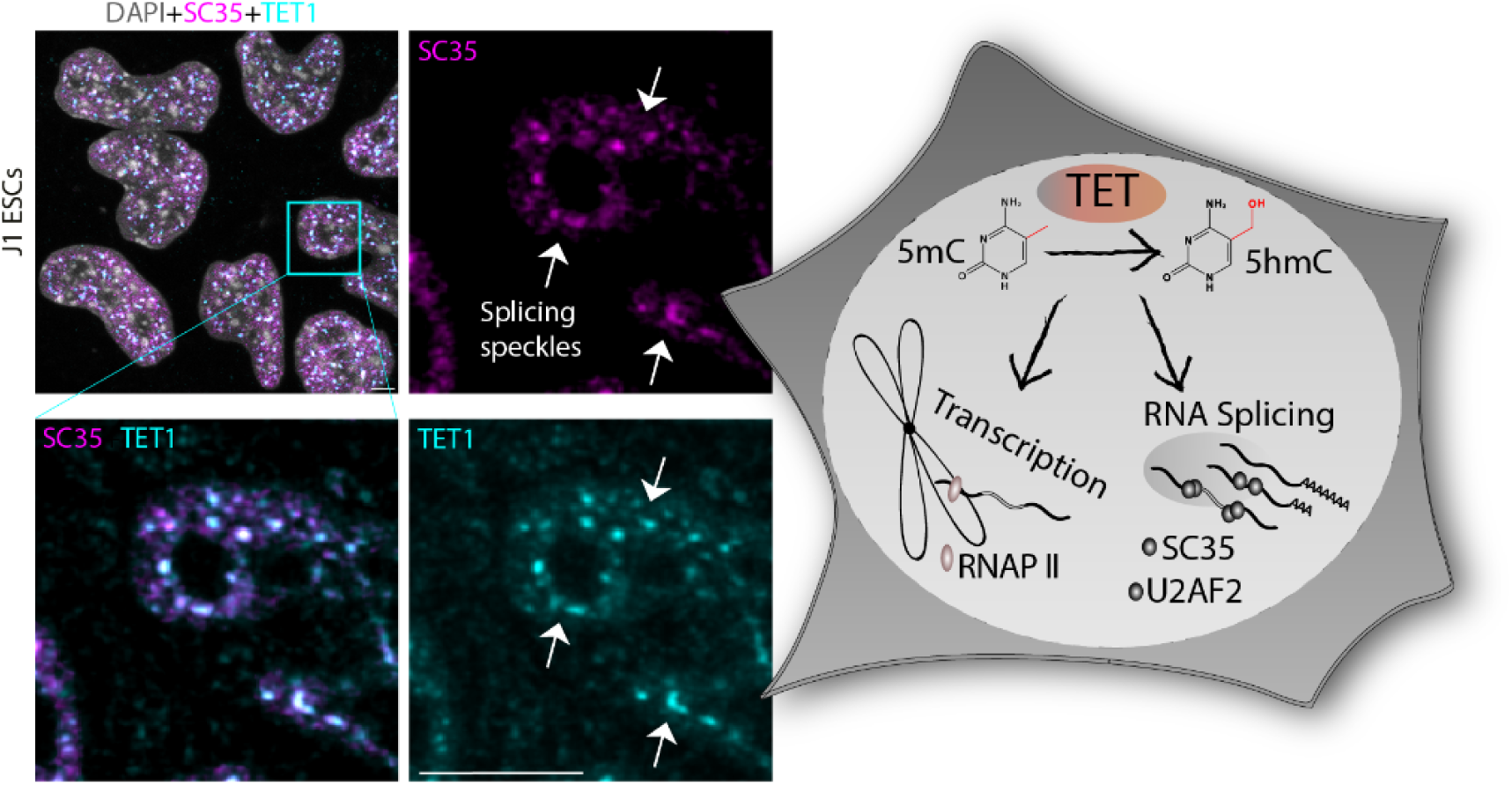

**Highlights:**

– TET1 localizes in splicing speckles in an RNA-dependent manner
– TET proteins, especially TET1, interact with the splicing factors U2AF1 and U2AF2
– TET proteins increase splicing efficiency, independent of their catalytic activity
– RNA 5-methylcytosine (m5C) oxidation to 5-hydroxymethylcytosine (hm5C) restores splicing efficiently in vitro

## Introduction

Cellular ribonucleic acids (RNAs) display many biological functions, and the RNA processing machinery is crucial for the cell. Protein-coding RNAs need to be processed to remove non-coding regions, a task performed by a macromolecular machine, the spliceosome. One of its components is SC35, belonging to the family of serine-arginine-rich proteins (SR proteins), and is involved in intron recognition and spliceosome assembly (Thomas et al., 2012). Its localization is mainly restricted to so-called nuclear speckles, also referred to as “SC35 domains” (Wansink et al., 1993), RNA–protein interchromatin granules found in the nucleoplasm of eukaryotic cells. The term “speckles” was first used in 1961, and two years later, these speckles were identified as RNA-containing interchromatin particles (Galganski et al., 2017). Speckles are enriched for RNA-binding proteins (RBPs) that belong mainly to the different subsets of pre-mRNA splicing factors (Perraud et al., 1979; Spector et al., 1991, 1983). Interestingly, SC35 distribution is unaffected by RNase A digestion, indicating that the association with speckles is based on protein-protein interactions rather than protein-RNA interactions (Spector et al., 1991). Besides the SR protein family, various other splicing factors contain SR domains. One of them is the U2 small ribonucleoprotein particle (snRNP) auxiliary factor 2 (U2AF2), with an N-terminal SR domain and three C-terminal RNA-binding domains (Carmo-Fonseca et al., 1992; Zamore et al., 1992). Its structure is conserved throughout different species, including *Drosophila melanogaster* (dU2AF50) and *Saccharomyces cerevisiae* (Mud2) (Fu, 1995). U2AF2 interacts with a smaller subunit called U2AF1, also containing an SR domain, which was found to interact with SC35 (Wu and Maniatis, 1993). During splicing, the freshly transcribed premature messenger RNA (pre-mRNA) undergoes a series of rearrangements, which can occur even during ongoing transcription. This is a fundamental process that increases the diversity of the transcriptome and, consequently, the proteome. The spliceosome complex plays a major role in its regulation and its importance is underlined by aberrant splicing events that were associated with different diseases, including diabetes, stroke, hypertension, and cancer (Bradley and Anczuków, 2023; Kashyap et al., 2022; Wang and Aifantis, 2020). Splicing is also related to biological processes like terminal erythropoiesis (Conboy, 2017) or neurogenesis (Fisher and Feng, 2022). Furthermore, alternative splicing events increase transcript diversity in different cell types and under variant conditions (Fisher and Feng, 2022). Internal base editing and splicing guarantee the exclusive continuance of coding sections, while distinct modifications at the 5′ and 3′ untranslated regions (UTR) ensure mRNA stability and correct subcellular localization (Martin and Ephrussi, 2009). Translational capacity is also dependent on RNA modifications like the incorporation of pseudouridine (Karikó et al., 2008) or N6-methyladenosine (m6A) (Mendel et al., 2021). Recently, it has been shown that splicing of mRNA is more efficient when tethered to the transcription elongation complex, as the newly synthesized RNA strand is extruded from RNA polymerase II (RNAPII) (Shenasa and Bentley, 2023).

In mammals, it is well established that DNA cytosine base modifications, especially 5mC, are important epigenetic marks and execute crucial functions in cellular processes (Trojer and Reinberg, 2007). 5mC is generated and maintained by DNA methyltransferases (MTases) called Dnmts (Bestor, 2000) and is mainly found in CpG dinucleotide-rich sites, resulting in dense packing of chromatin (Bird, 2002). In 1978, m5C was also detected in the rRNA of wheat seedlings (Rácz et al., 1978), raising the question of whether it can fulfill regulatory functions in RNA. Since then, it has been shown that m5C contributes to the structural stabilization of RNA molecules impairing the recognition of endogenous RNAs by the innate immune system (Motorin et al., 2010). In lncRNAs, m5C can influence their binding dynamics to chromatin-modifying complexes. Furthermore, it can take part in the generation of individual sets of mRNAs by affecting the metabolism of microRNAs (L. C. Huber et al., 2015). In tRNA and rRNA, m5C has been extensively studied, but little is known about its role in mRNA, in part, due to the lack of effective separation and purification technologies (Guo et al., 2021). Similar to DNA, methylation of cytosines in RNA is catalyzed by specific RNA MTases (RMTases) like TRDMT1 and the nucleolar protein 1 (NOL1)/NSUN protein family (Goll et al., 2006; Reid et al., 1999). These proteins use S-adenosylmethionine (SAM) as a methyl group donor (Motorin et al., 2010). However, the identity of m5C modifiers and erasers is still vague. Since the discovery of m5C, more than 300 RNA modifications have been identified (Arzumanian et al., 2022; Cappannini et al., 2024), N6-methyladenosine (m6A) being one of the most relevant. This modification was shown to be the substrate for iterative oxidation to N6–hydroxymethyl adenosine and further to N6-formyladenosine, in a reaction catalyzed by the AlkB protein family of dioxygenases (Fu et al., 2013; Jia et al., 2011). Notably, members of the AlkB family and the DNA modifying ten-eleven-translocation (TET) family share structural similarities. TET proteins are Fe(II) and 2-oxoglutarate-dependent dioxygenases and are known to act on DNA by oxidizing 5-methylcytosine (5mC) to 5-hydroxymethylcytosine (5hmC) and further to 5-formylcytosine (5fC) and 5-carboxylcytosine (5caC). The mammalian TET family consists of three members, TET1, TET2, and TET3, sharing a conserved catalytic domain at the C-terminus (Hu et al., 2013). Interestingly, TET1 is the most ubiquitously expressed member across different tissues (Ito et al., 2011). The AlkB family of dioxygenases can use DNA and RNA as substrates (Falnes et al., 2004; Fedeles et al., 2015); however, whether the related TET proteins do the same was unclear. Later on, it was shown that RNA can be the substrate of TET dioxygenases, as the occurrence of RNA 5-hydroxymethylcytosine, termed hm5C, is dependent on the presence of TET proteins (Fu et al., 2014). Furthermore, hm5C has been detected in different tissues, amongst others, the brain, heart, pancreas, and spleen. The iterative oxidation of m5C to hm5C in total RNA was demonstrated in a mouse model, suggesting a conserved process that could have critical regulatory functions inside cells (S. M. Huber et al., 2015). In addition, 5-formylcytidine (f5C) was detected in mammalian RNA, where its formation is currently assumed to be catalyzed by TET proteins (Zhang et al., 2016). Interestingly, TET has been shown to interact with microRNAs regulating hm5C in the adult brain (Kremer et al., 2018), and hm5C was found in the RNA of Drosophila S2 cells (Delatte et al., 2016). These cells contain a conserved TET analog (called dTET), which favors the conversion of m5C to hm5C in polyadenylated RNA, resulting in a more efficient translation of the hydroxymethylated mRNA.

Although the function of TET proteins in the oxidation of m5C seems clear, a lot about the role of RNA modifications and their effect on RNA metabolism remains elusive. Splicing has emerged as a key regulator of neural development, where spliceosome dysfunction causes a series of neurodevelopmental disorders with similar features (D. Li et al., 2024). In this context, investigating the so-called “splicing code” and novel factors involved in splicing regulation has gained interest. In this study, we address the connection between RNA splicing and TET dioxygenases and their effect on RNA splicing. We show that TET proteins localize to splicing speckles in mouse, human, and Drosophila cells and tissues. This association with speckles is RNA-dependent, as is their interaction with splicing factors underlined. Furthermore, we show that m5C blocks splicing, while its oxidative derivative, hm5C, restores splicing activity. Overall, our results provide novel insights into how RNA splicing is regulated, with TET proteins and their oxidation product hm5C having an impact on it.

## Results and discussion

### TET proteins interact with the splicing factors U2AF1 and U2AF2

In the course of mass spectrometry experiments investigating TET proteins interactome and posttranslational modifications (Bauer et al., 2015), we found a significant enrichment of splicing factors in the proteins detected by LC-MS/MS. In this analysis, GFP-tagged murine TET proteins or GFP as control (Supplementary Table S1) were expressed in human HEK293T cells. GFP-tagged proteins were pulled down from whole cell lysates with GFP-binder beads and subjected to LC-MS/MS (Fig. 1). An overall sequence coverage of ∼50% was achieved for TET1, ∼60% for TET2, and ∼65% for TET3 (Supplementary Data S1). Volcano plots show the analysis of these experiments for TET1 (Fig. 1A), TET2 (Fig. 1B), and TET3 (Fig. 1C), where each circle/triangle represents a protein detected by LC-MS/MS and the X-axis depicts differences in protein abundance in the respective pulldowns as Log2 fold change (GFP-TET versus GFP control). Factors more enriched in TET-IP (immunoprecipitation) have positive Log2 fold change values (label-free quantification by MaxQuant). Mass spectrometry analysis of TET-associated factors showed stronger enrichment for the splicing factors U2AF1, U2AF2, and SRSF2 (also known as SC35) (blue triangles) for TET1 pull-downs, while this enrichment was less significant for TET2 and TET3. The Venn diagram in Fig. 1D shows splicing-associated factors that were found to interact with all three TET proteins and illustrates the overlap of splicing factors identified in all TET pull-downs. Again, TET3 and TET2 showed less enrichment of splicing-associated factors (6 and 5 respectively) compared to TET1 (11 splicing factors). Nonetheless, there was a shared set of splicing factors interacting with all three proteins. Specifically, the proteins U2AF1, U2AF2, and SC35 were significantly enriched in the TET1 and TET2 pull-down. These proteins are critical components of the spliceosome and involved in pre-mRNA processing, which suggests a functional interaction between TET dioxygenase enzymes and splicing machinery. Given the role of TET proteins in DNA demethylation and potential involvement in RNA modifications (e.g., hm5C in RNA), these interactions could reflect a regulatory role in gene expression via splicing.

**Figure 1.**
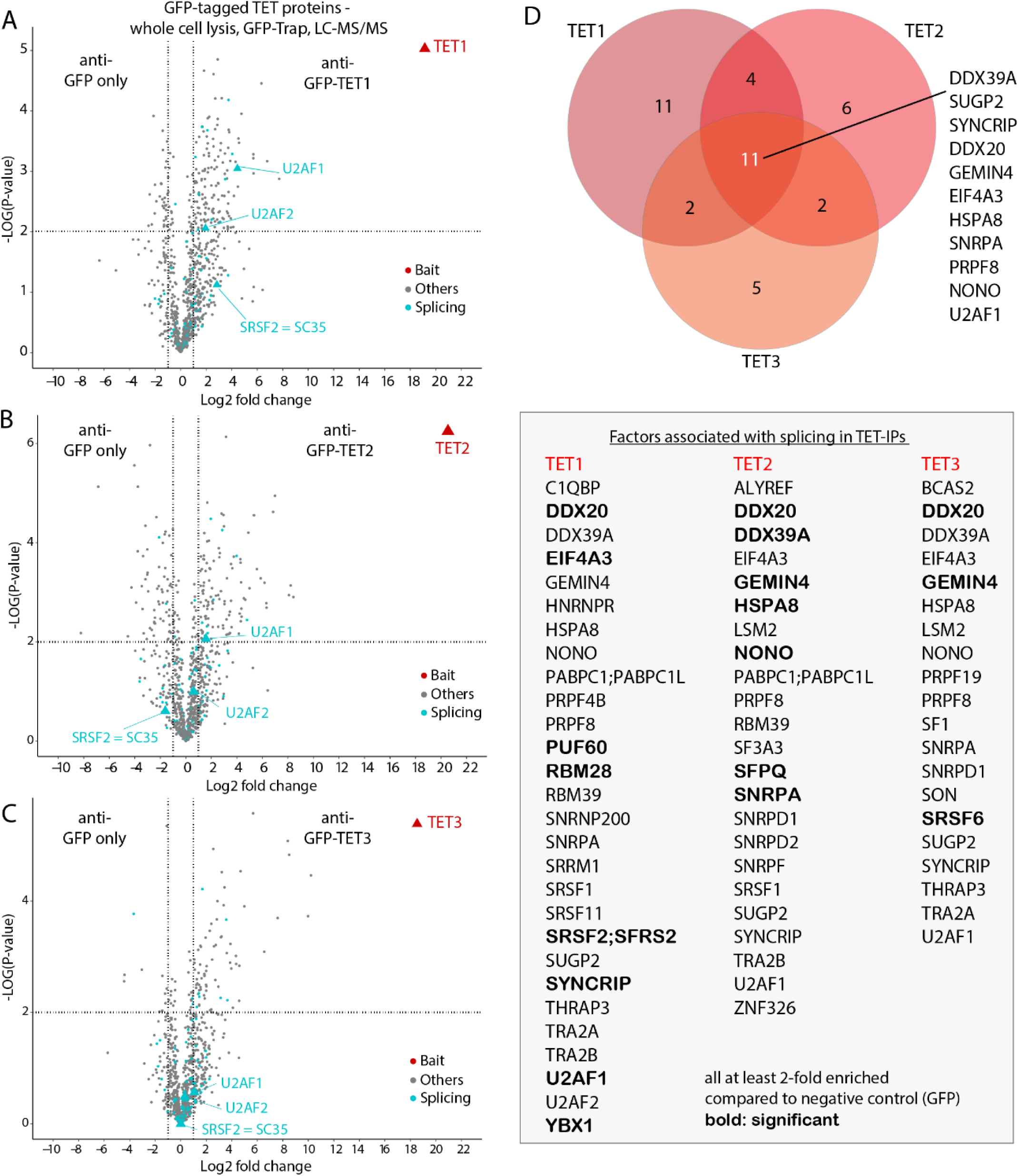
Mass spectrometry analysis reveals novel interactions between TET proteins and splicing factors. **(A)** TET1, **(B)** TET2, and **(C)** TET3 mass spectrometry analysis of TET-associated factors. GFP-tagged murine TET proteins or GFP as control were expressed in human HEK293T cells. GFP-tagged proteins were pulled down from whole cell lysates with the GFP-binder beads and subjected to LC-MS/MS. Volcano plots: Each circle/triangle represents a protein detected by LC-MS/MS; X-axis depicts differences in protein abundance in the respective pulldowns as log2 fold change (GFP-TET versus GFP control); factors more enriched in TET-IP have positive Log2 fold change values (label-free quantification by MaxQuant) Y-axis depicts the negative log10 of the p-value of a student’s t-test (triplicate samples) dashed black lines: significance border (FDR=0.05, S0=2), blue circles/triangles denote splicing factors. A higher value corresponds to a lower p-value, indicating stronger statistical significance. Proteins outside the dashed lines (upper right or left corners) are significantly enriched or depleted, respectively. **(D)** Venn diagram shows splicing-associated factors that were found to interact with different TET proteins. TET3 and TET2 showed slightly less enrichment of splicing-associated factors compared to TET1, as shown in the list below.

Mass spectrometry studies provided insights into both shared and specific protein interactions for TET proteins, suggesting that a core set of splicing factors may be associated with TETs, while unique interactions may point to specific functions for each TET protein in different cellular contexts. Spliceosome assembly occurs by ordered interaction of the spliceosomal snRNPs (small nuclear ribonucleoproteins) and numerous other splicing factors. During exon definition, the U1 snRNP binds to the 5′ss downstream of an exon and promotes the association of U2AF (U2AF1 and U2AF2) with the polypyrimidine tract/3′ss upstream of it. Splicing enhancer sequences within the exon (ESEs) recruit proteins of the SR protein family (among them SC35), which establish a network of protein-protein interactions between the different components of the spliceosome (Fig. 2A) (Will and Lührmann, 2011). To gain more insights into the role of TET proteins in this process, we performed an in silico screening for protein-protein-RNA interactions between TET and the splicing factors U2AF1, U2AF2, and SC35 using AlphaFold3 (https://alphafoldserver.com/) (Abramson et al., 2024). We further expanded this modelling by incorporating a short RNA sequence within the intron/exon limit. Previous studies reported significant decreases in the sensitivity of AlphaFold for complex structure predictions when using full-length sequences (Lee et al., 2024). Therefore, after full-length protein predictions, we also designed and paired protein fragments either consisting of individual folded or disordered regions in TET proteins and U2AF1, U2AF2, or SC35 subunits. These fragments were chosen based on the PAE (predicted aligned error) values for each pair of residues in the full-length predictions, selecting the protein domains predicted to interact with a low error rate. For most of the structures, these fragments included the cysteine-rich domain in the catalytic domain of TET proteins and the RRM (RNA recognition motif) for the three splicing factors. A diagram of the proteins including the most relevant domains and the fragments used for AlphaFold predictions is shown in Fig. 2B. In total, we conducted 121 AlphaFold predictions in our screening for all TET proteins in combination with the different splicing factors and between U2AF and SC35 proteins, all in the presence of the short RNA fragment. The heatmap in Fig. 2C shows the ipTMs (interface-predicted template modeling) scores for all these predictions. The ipTM and pTM (predicted template modeling) scores measure the confidence of the overall structure (shown in Fig. S1 and Supplementary Data S2). The pTM is an integrated measure of the prediction, showing the accuracy of the predicted structure for the full complex as a score for the superposition between the predicted structure and the hypothetical true structure. However, the ipTM measures the accuracy of the predicted relative positions of the subunits within the complex, or how they are predicted to interact, being a metric of the predicted interface of the interaction. Therefore, we ranked the resulting models by model confidence using both ipTM and pTM scores. The highest confidence models were selected for each TET protein, with an ipTM of 0.68 for the interaction between TET1_aa1094-1736 and U2AF2_2RRM (Fig. 2D), ipTM of 0.54 for TET2 and U2AF1 (Fig. 2E), and ipTM of 0.57 for TET3_aa560-1170 and U2AF2 (Fig. 2F). The three structural models comprising TET (green), the respective splicing factor (cyan), and RNA (pink) show the layout of the interaction from different angles, with multiple points of predicted interaction between both proteins and the RNA. For better visualization, magnifications of these proximal regions show the U2AF structure and exclusively the residues of TETs located within a range of 5 angstroms (Å). Additional images of the structures and the residue interactions within a range of 4 angstroms (yellow dashed lines) are shown in Fig. S1B-D. Interestingly, a short fragment of U2AF2 (aa130-143) was predicted to interact with TET proteins with the highest scores (0.74 for TET1_aa1409-1686) (Fig. S1B). This fragment, which corresponds to 13 residues, is located in proximity to the second RRM domain in U2AF2 (2RRM) and is not present in U2AF1 or SC35. The results of the interactions screening and structural modeling show that TET1 is predicted to interact mostly with U2AF2, while TET2 and TET3 interactions show higher confidence scores for U2AF1. On the other hand, only TET3 showed an ipTM of 0.7 in the interaction with SC35 but with a lower pTM of 0.59. Higher-ranked predictions were validated using AlphaFold2 (Multimer), obtaining similar scores trends and structural models. Overall, AlphaFold screening supports our initial finding from mass spectrometry, illustrating the structural interplay between TET proteins, splicing factors, and RNA, while also confirming potential differences between the different TET proteins.

**Figure 2.**
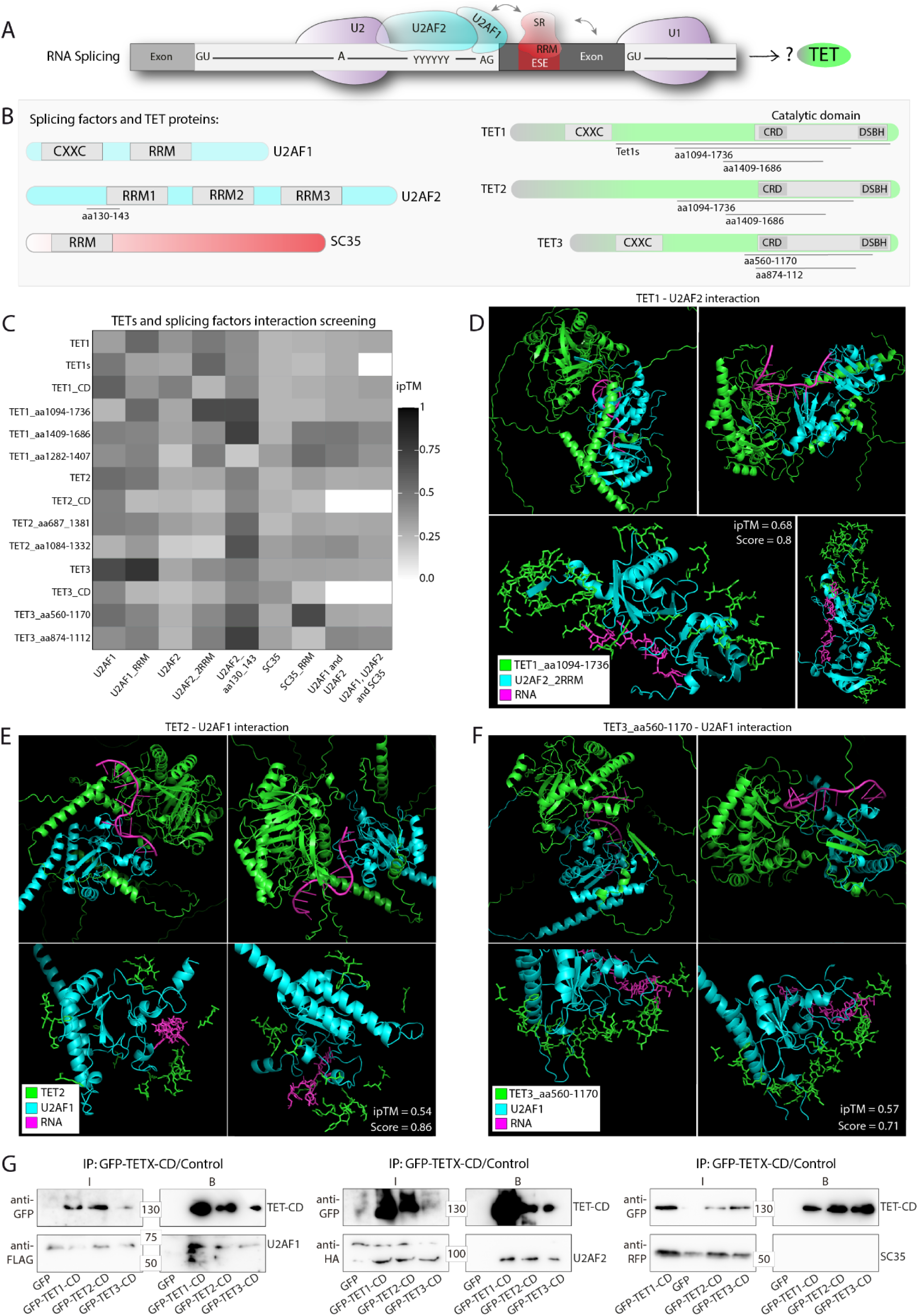
Screening and validation of TET protein-protein-RNA interactions by structural modeling with AlphaFold 3. **(A)** The diagram shows the process of RNA splicing and the proteins involved, with a pre–mRNA harboring exons and introns (light grey) which are removed during splicing. The splice sites at the start “GU” and the end “AG” of the intron are shown, together with the branch point “A”. The polypyrimidine tract located upstream of the 3’ splice site is represented as “YYYYY”. U1 and U2 snRNPs, components of the spliceosome, are responsible for recognizing splicing sites. U2AF1 and U2AF2 proteins assist in binding to the 3’ splice sites and branch points. SR proteins (like SC35) bind to exonic splicing enhancers (ESE) to enhance splicing accuracy. **(B)** Schematic representation of U2AF1, U2AF2, and SC35 harboring different RRM domains (RNA recognition motifs), and TET proteins, with their domains, represented: CXXC (zinc finger domain), the conserved catalytic domain in the C-terminus, including the cysteine-rich domain (CRD) and the double strand beta-helix domain (DSBH), with an insert in between. In addition to the full protein sequences, different fragments of the aminoacid sequences and domains were used to predict protein-protein-RNA interaction, as indicated in the scheme. **(C)** Heatmap showing the ipTM score (interface predicted Template Modeling score) obtained for AF structure models in the screening for all interactions tested between TET proteins and the splicing factors U2AF1, U2AF2, and SC35. Labels on the x and y axis indicate the paired protein fragments for structural modeling, including RNA in the prediction. White tiles indicate pairs that were not subjected to structural modeling. **(D)** The highest-scored structural models were obtained for TET1-U2AF2-RNA, **(E)** TET2-U2AF1-RNA, and **(F)** TET3-U2AF predictions. The full structural models are shown from different perspectives (top part), and magnifications are shown at the bottom. In the magnification, only residues within a maximum distance of 5 angstroms to U2AF are shown. **(G)** Co-immunoprecipitation analysis of TET proteins interaction with splicing factors. The catalytic domain (CD) of TET1, TET2, and TET3 or GFP as control were co-immunoprecipitated using GFP-binder beads. Pull-down fractions were analyzed by western blot using anti-FLAG antibody (left, U2AF1), anti-HA antibody (center, U2AF2), or anti-RFP (right, SC35). Inputs “I” and bound “B” fractions are shown. Full blots images and replicates are shown in Figure S2.

Next, we validated AlphaFold3 predictions between TET proteins and splicing factors by co-immunoprecipitation (Fig. 2G). To this end, HEK-EBNA cells were transiently co-transfected with plasmids coding for GFP-tagged TET1, TET2, or TET3 catalytic domains (CD) or GFP as control, and U2AF1-FLAG, U2AF2-HA, or mCherry-SC35. GFP-tagged proteins were immunoprecipitated using a GFP-binder (Rothbauer et al., 2008), and cell lysates were analyzed by western blot. We found that all TET proteins were able to pull down U2AF1 and U2AF2 but not SC35 (Fig. 2G and Fig. S2A-B). Additionally, we co-transfected cells with YFP-SC35 and mCherry-tagged TET1, TET2, or TET3 catalytic domains, immunoprecipitated YFP-SC35 as before, and analyzed cell lysates by western blotting. In line with the reverse experiment, immunoprecipitated SC35 was not able to pull down either of the TET proteins (Fig. S2B-right) (Full blot images and replicates are shown in Fig. S2). This again confirmed our AlphaFold protein-protein interaction predictions, showing that U2AF1 and U2AF2 proteins form a multimer complex (Wu and Maniatis, 1993) that interacts directly with the catalytic domain of TET proteins. However, there is no direct interaction between SC35 and TET proteins. These findings support the initial hypothesis based on mass spectrometry data and AlphaFold screening, showing that TETt1, TET2, and TET3 interact with the splicing factors U2AF, implicating a potential role in the regulation of RNA splicing.

### TET proteins localize at splicing speckles in mouse, human, and Drosophila

Previous studies have shown that RNA serves as a substrate for TET proteins and that hm5C levels increase upon TET overexpression (Fu et al., 2014). This evidence supports the role of TET DNA dioxygenases in regulating RNA modifications, which is consistent with our finding that TETs interact with splicing factors. Therefore, we investigated the connection between TET proteins and RNA splicing, investigating whether TET proteins localize to splicing speckles. For these experiments, we focused mostly on TET1 since it showed the highest interaction score with U2AF2 and it is abundantly distributed in a higher number of cell lines and tissues (Santiago et al., 2014). The splicing factor SC35 is a reliable marker for these subnuclear compartments, and specific antibodies are available. Using immunofluorescence, we addressed a potential colocalization of TET in splicing speckles. We performed immunostainings for both TET1 and SC35 proteins in J1 mouse embryonic stem cells (mESCs) versus differentiated cells (Fig. 3A). The localization of TET1 and SC35 in splicing speckles was observed in mESCs and across various differentiation states: embryoid bodies (EB) derived from J1 ESCs, mouse embryonic fibroblasts (MEF), and mouse tail fibroblasts (MTF), using Oct4 as a pluripotency marker. To validate this observation, colocalization between TET1 and SC35 (grey) and colocalization between TET1 and DAPI (black) were quantified using the H-coefficient (Fig. 3A-right). Here, positive colocalization was found for TET1 and SC35 (H-coefficient higher than 1) in all cell lines. However, anti-colocalization (negative values) was found for TET1 and DNA-dense regions stained with DAPI, which are devoid of splicing speckles. From these experiments, we can conclude that TET1 proteins colocalize with splicing speckles in different cell lines and differentiation stages. In addition, similar observations were found performing immunofluorescence for TET2 and TET3 in MTFs (Fig. S3) and human fibroblast Bj-hTERT (Fig. S4), where all three TET proteins were observed to colocalize with SC35 in splicing speckles. Localization of TET1 in the nucleoplasm also agrees with the observation of the Human Protein Atlas (https://www.proteinatlas.org/) in different human cell lines. Next, we performed immunostainings in MTFs transiently transfected with tagged U2AF proteins (U2AF1-Flag and U2AF2-HA) (Fig. 3B). MTF cells were chosen for these experiments because they show high endogenous TET1 levels. The quantification of TET1 and U2AF colocalization in speckles also showed H-coefficient values higher than 1 for TET1 and both U2AF proteins and values close to 0 (no colocalization) for dense DNA regions (DAPI-rich) (Fig. 3B-right). This confirmed the subnuclear association of TET1 proteins with RNA splicing speckles and splicing factors.

**Figure 3.**
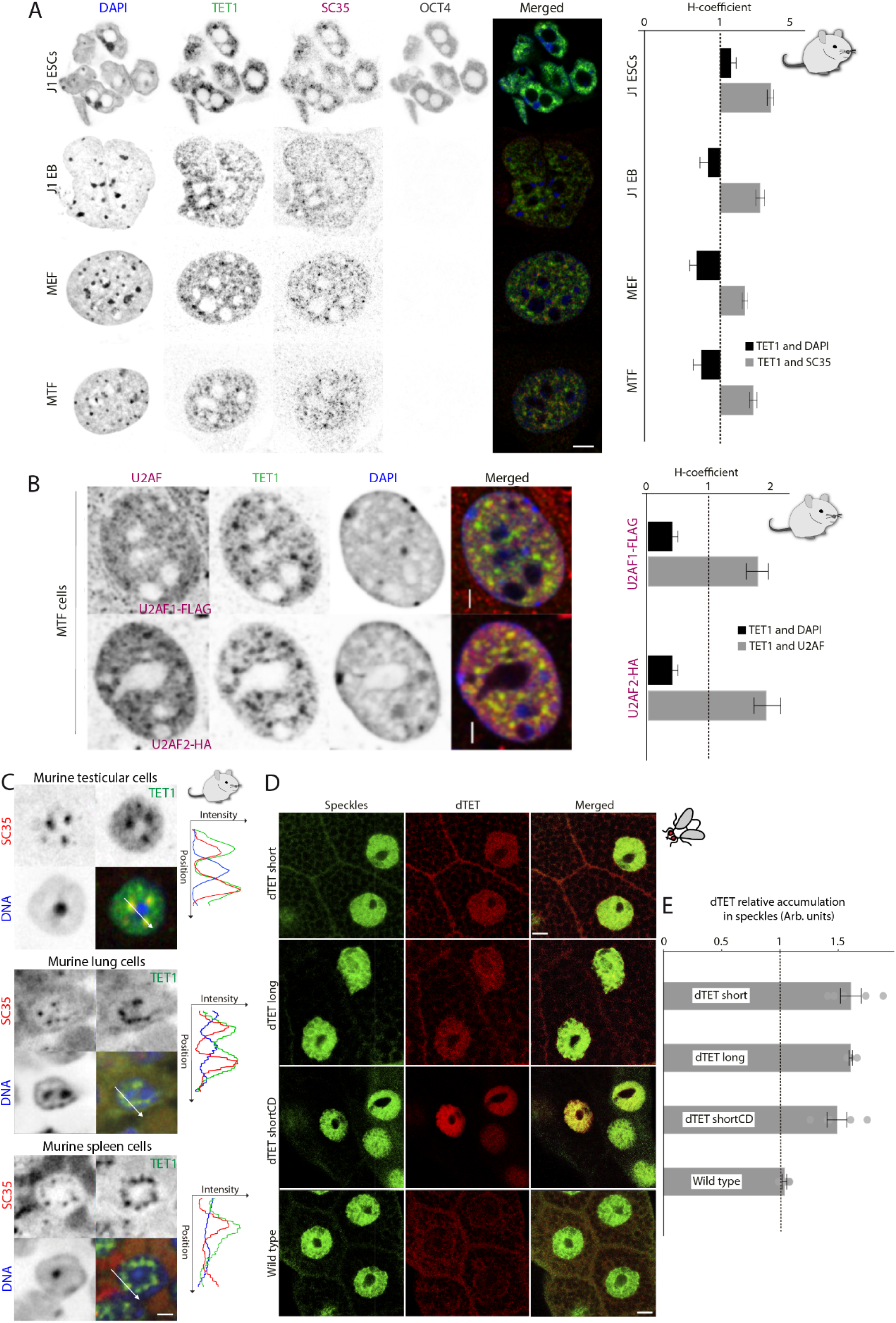
TET1 localizes at splicing speckles, colocalizing with splicing factors in mouse and Drosophila tissues. **(A)** Immunostaining of TET1 and SC35 in various differentiated mouse cell lines with quantified colocalization represented by the H-coefficient. J1 ESCs, embryoid bodies (EB) derived from J1 ESCs, mouse embryonic fibroblasts (MEF), and mouse tail fibroblasts (MTF) were stained for TET1 (green) and SC35 (red). DNA was counterstained with DAPI (blue). To distinguish the undifferentiated state, Oct4 was used as a pluripotency marker. The Oct4 channel was omitted for the merge. Colocalization between TET1 and DAPI (black) as well as TET1 and SC35 (dark grey) was quantified using the H coefficient (n=10 cells) and is shown as a barplot on the right-hand side. **(B)** MTF cells were immunostained for tagged U2AF proteins and endogenous TET1 (green). Ectopically expressed U2AF1-FLAG or U2AF2-HA (red) were visualized using anti-HA or anti-FLAG antibodies. DNA was counterstained with DAPI (blue). The barplot illustrates the H-coefficient (n=10) of TET1 and DNA (black) or U2AF proteins (grey). Scale bar: 5 µm. **(C)** TET1 localizes to SC35-positive speckles in vivo. Paraffin sections of murine testis, lung, and spleen were immunostained with antibodies against TET1 and SC35. DNA was counterstained with DAPI. Line-profile analysis of selected regions denote TET1 colocalization with SC35 and anti-colocalization with DNA. **(D)** Colocalization of dTET and nuclear speckles in Drosophila salivary gland cells. Different HA-tagged dTET isoforms (short (row 1), long (row 2), and catalytically dead (cat. dead) short isoform (row 3) were expressed from UAS inserts by elav^C155^GAL4, and compared with wild type (raw 4). Third instar salivary glands were stained for speckles with anti-SC35 antibodies and dTET with anti-HA antibodies. The overlay of dTET and SC35 are shown in the merge. The boxplot on the right shows the quantification by image analysis of dTET accumulation in speckles. Values exceeding 1 (dashed black line) denote colocalization with SC35. Scale bar: 15 µm. Error bars represent the standard deviation.

To investigate the nuclear localization of endogenous TET1 in splicing speckles in vivo, we performed TET1 and SC35 immunostainings in mouse tissue sections. The most significant TET1 signals were observed in the testes sections, lung, and spleen (Fig. 3C, images of the sections in Figure S5, S6, and S7). TET1 was detected in low-density DNA regions in these immunostainings, clearly colocalizing with SC35-positive splicing speckles. These results show the subnuclear localization of endogenous TET1 proteins at splicing speckles in vivo, pointing to a physiological role of TET1 proteins regulating gene expression by alternative splicing. On the one hand, TET1 could play a role in chromatin reorganization and decondensation by erasing 5mC, which keeps chromatin in a densely packed state (Razin and Cedar, 1977). In line with this hypothesis, TET1s (short isoform) has been shown to cause 5hmC increase and decondensation of heterochromatic regions upon recruitment (Arroyo et al., 2022). On the other hand, TET1 may regulate splicing, leading to the formation of cell-type-specific proteins in chromatin reorganization (Kanto et al., 2016). TET1 localization with SC35 at nuclear speckles may have several physiological consequences. Accordingly, previous studies detected hm5C in RNA of different tissues, like the heart and spleen (Fu et al., 2014). These studies, together with our previous results showing TET1 interaction with splicing factors, lead to the prospective role of TET proteins regulating splicing, either via their catalytic activity and m5C oxidation, interaction with splicing factors and/or non-catalytic functions, some of them discovered recently (Arroyo et al., 2022; Chrysanthou et al., 2022; Ketchum et al., 2024).

To investigate whether TET’s association with splicing speckles was related to DNA or RNA modifications (5mC versus m5C), we selected Drosophila melanogaster as a model. In human and mouse cells, it is not possible to distinguish whether this association is due to TET activity on methylated DNA or RNA. However, previous studies have shown that DNA methylation in Drosophila is nearly undetectable (Capuano et al., 2014; Dunwell and Pfeifer, 2014; Krauss and Reuter, 2011), while RNA methylation-associated processes are present and functionally important. Drosophila has only one TET gene, which is alternatively spliced to generate two isoforms. While the long isoform with a CXXC zinc-finger DNA binding motif is similar to vertebrate TET1 and TET3, the short isoform without this motif resembles vertebrate TET2 (Wang et al., 2018). We have shown that human and mouse TET proteins localize to speckles (Fig. 3A-C, Fig. S3-S7), which are sites for pre-mRNA splicing (Maniatis and Reed, 2002). Next, we investigated TET subnuclear localization in Drosophila to clarify whether it is DNA methylation-dependent. For this purpose, we used Drosophila salivary glands expressing different TET isoforms (dTET short and dTET long), including a catalytically-dead short isoform (dTET shortCD), using C-terminally HA-tagged UAS (Upstream Activation Sequence) constructs expressed with elavC155GAL4. In the large Drosophila salivary gland cells, speckles can also be visualized with antibodies against human SRSF2 (SC35) (Allemand et al., 2001; Prasanth et al., 2000). Interestingly, by performing immunofluorescence analysis we found that all three dTETs colocalize with speckles in the nucleus but were not detected in the nucleolus and that this localization is not dependent on the catalytic activity of dTET (Fig. 3D). Then, to analyze whether loss of TET affects alternative splicing in Drosophila, we analyzed a differential gene expression RNA-seq data set for changes in splicing, finding 594 and 326 two-fold up- or down-regulated genes, respectively (Supplementary Data S3) (Boulet et al., 2023). This analysis revealed significant changes (>30% changes and p<0.01) on exon skipping in 27 genes (Fig. 4A), intron retention in 16 genes (Fig. 4B), of 5’ and 3’ splice sites for 42 and 19 genes, respectively (Fig. 4C-D), and 26 mutually exclusively spliced exons in 7 genes (Supplementary Data S4). Among changes in the genes with mutually exclusive spliced exons, we found several events for the Dscam1 gene. In Drosophila, this is one of the most complex alternatively spliced genes harbouring three variable clusters, exon 4, exon 6, and exon 9, with 12, 48, and 33 variables, respectively (Hemani and Soller, 2012). Furthermore, gene ontology (GO) analysis of differentially expressed genes revealed a significant number of genes involved in detoxification including several Cytochrome C genes and metabolic genes (Fig. 4E). GO analysis for changes focused on alternative 5’ splice sites revealed a significant enrichment (p<0.05) in neurodevelopmental genes, which is consistent with the defects in TETnull mutants (Delatte et al., 2016; Ismail et al., 2019).

**Figure 4.**
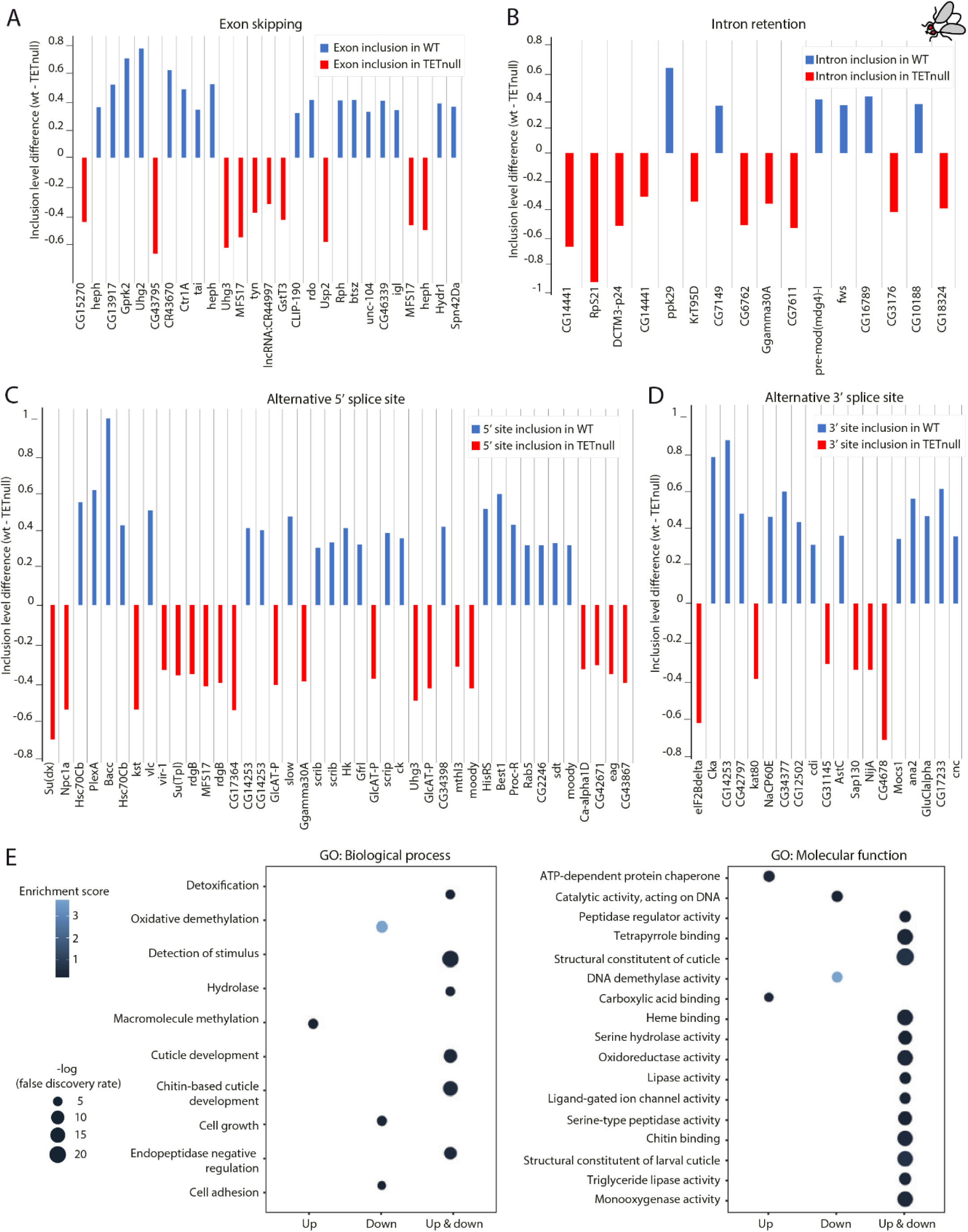
RNA-seq analysis of alternative splicing in Drosophila wild-type versus TETnull. **(A)** Barplot showing the inclusion level difference of exon skipping genes between wild-type (wt) and TETnull samples. **(B)** Barplot showing the inclusion level difference of intron retention between wild-type (wt) and TETnull samples. **(C)** Barplot showing the inclusion level difference of alternative 5’ splice sites between wild-type (wt) and TETnull samples. **(D)** Barplot showing the inclusion level difference of alternative 3’ splice sites between wild-type (wt) and TETnull samples. For (A), (B), (C), and (D), genes showing significant changes are shown (>30% changes and p<0.01, Supplementary Data S4), ordered from lowest to highest p-value from left to right. Inclusion level measures the frequency of a particular event (exon skipping, intron retention, 5’ or 3’ alternative splicing sites) in the final mRNA transcript, represented as a value between 0 (always skipped) and 1 (always included). Inclusion level difference is calculated as the inclusion level in wild-type minus inclusion level in TETnull condition. Blue bars with positive values indicate an increased inclusion in wild type, while red bars with negative values indicate an increased inclusion in TETnull. **(E)** Gene ontology (GO) analysis of differentially expressed genes (Supplementary Data S3) are shown on the left for biological process and on the right for molecular function, with enrichment indicated in blue color and -log(false discovery rate) by circle size.

Splicing speckles localize in the proximity of transcription sites and form subcellular domains called gene expression factories (Maniatis and Reed, 2002). Since Drosophila lacks 5mC methylation in DNA (Raddatz et al., 2013) but still has a TET gene highly homologous to the vertebrate TET proteins, we hypothesized a DNA methylation-independent function of TET proteins in chromatin-associated regulation of transcription. Interestingly, we found a colocalization of TET with elongating RNA Polymerase II (RNAPII) in Drosophila salivary glands (Fig. S8A and Fig. S8B). TET localization at splicing speckles and TET colocalization with RNAPII were found in all conditions in Drosophila cells. The same colocalization between TET1 and RNAPII was found in mouse J1 mESCs, in addition to TET1 localization at splicing speckles (Fig. S8C-D). These results show that both elongating RNAPII and TET colocalize, with TET located at splicing speckles in the so-called “gene expression factories”. Therefore, TET proteins may have a conserved role in regulating mRNA splicing and gene expression by alternative splicing. Interestingly, the RNAPII CTD has been shown to coordinate pre-mRNA processing events (de Almeida and Carmo-Fonseca, 2008).

### TET1 association with speckles and splicing factors interaction is RNA-dependent

To study the characteristics of the association of TET1 with splicing speckles and whether it is dependent on the interaction with splicing factors or with RNA, we performed live-cell RNase extraction followed by immunostaining in mouse tail fibroblast (MTF) cells. Representative images obtained by confocal microscopy are shown in Fig. 5A, illustrating the total reduction of TET1 signal in extracted cells treated with RNase. In addition, the quantification of propidium iodide staining (which stains both DNA and RNA) shows a clear reduction after RNase treatment and extraction, corresponding to the RNA being washed out (Fig. 5A-right). Therefore, the association of TET1 with splicing speckles is completely lost after RNA removal, while speckles are still visible based on the SC35 signal. This suggests that TET1’s association with splicing speckles is primarily RNA-dependent. Therefore, RNA presence is a prerequisite for the reported colocalization and protein-protein interaction with splicing factors like U2AF. To verify this observation, we performed a fluorescent three-hybrid assay (Duan et al., 2021) in live cells to test the direct interaction of TET1-CD with an mRNA mimic. In this essay, the mRNA mimic (ms2-PABPC1-mCherry) is cotransfected in BHK cells (Tsukamoto et al., 2000) together with an RNA trap (MCP-EGFP-LacI), and the protein of interest, in this case PABPC1-mCherry (known to interact with the mRNA mimic) (Duan et al., 2021) or mCherry-TET1-CD. The RNA trap is tethered to the LacO locus in this cell line, which is visible as a big focus due to the EGFP fluorescence. This mRNA mimic is trapped at the LacO locus, and in the case of RNA-protein interaction, the recruitment of the target proteins by the mRNA mimic takes place. When this occurs, mCherry accumulation at the LacO can be imaged and quantified by live cell confocal microscopy (Fig. 5B). After image analysis, we found a significant increase in the relative accumulation of TET1-CD in the LacO, as a result of its interaction with the mRNA mimic, compared to the negative control (without the mRNA and showing no mCherry accumulation at all), and with the positive control also with PABPC1-mCherry accumulation (Fig. 5C-D). Thus, TET1 can be recruited by an mRNA mimic in live cells, confirming that TET1’s association with speckles is RNA-driven. In addition, we investigated the effect of RNA removal in the AlphaFold structural models, selecting the highest-scored models from Fig. 2. As shown in the heatmaps of Fig. 5E, almost all ipTMs and pTMs scores of previous predictions dropped to much lower values after removal of the short RNA sequence, demonstrating that RNA presence plays a crucial role in the interaction between TET and splicing factors (Supplementary Data S2). Structural modeling also showed that the TET1 catalytic domain alone is predicted to interact with RNA, and this interaction displays similar confidence scores to AlphaFold3 structural predictions for TET1-DNA interaction (Fig. 5F and Fig. S9).

**Figure 5.**
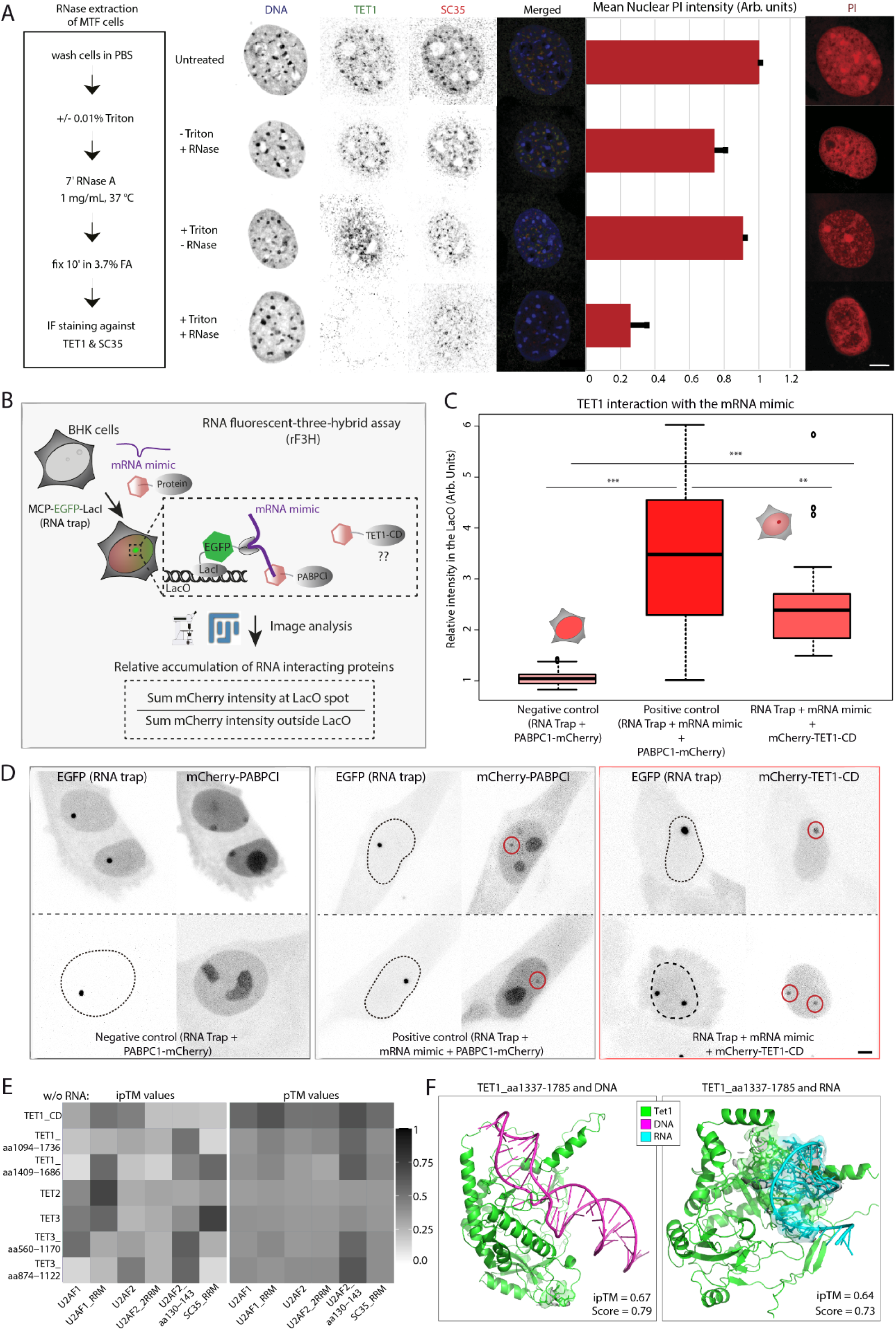
TET1 associates with RNA in splicing speckles and is recruited by an mRNA mimic in vivo. **(A)** Live cell RNase extraction of MTF cells. On the left, the scheme shows the experimental pipeline to study TET1 speckle association upon RNase A treatment. Mouse tail fibroblasts were seeded on gelatine-treated glass coverslips and 24 hours later pre-extracted with 0.01% Triton in PBS, followed by incubation with 1 mg/mL RNase A for seven minutes. Afterward, cells were fixed with paraformaldehyde and immunostained with antibodies against TET1 and SC35. Representative confocal microscopy images are shown in the center. As treatment control, similarly treated cells were stained with propidium iodide (PI), and the respective signal was quantified by high-content microscopy (n>1000 cells). Representative images of PI staining are shown on the right. Scale bar = 5 µm. **(B)** TET1-CD is recruited with a mRNA mimic in live-cell rF3H assay (RNA fluorescent three-hybrid assay). Scheme of the rF3H assay. Briefly, genetically modified BHK cells with many copies of the LacO were transfected with an RNA trap, an mRNA mimic, and PABPCI-mCherry or mCherry-TET1-CD. The RNA trap is fused to LacI (for LacO targeting) and EGFP (for visualization of the LacO locus). After live-cell imaging of transfected cells, the interaction between the mRNA mimic and the mCherry proteins is quantified by image analysis. **(C)** Boxplot showing the quantification of the relative accumulation of mCherry proteins at the LacO. Two independent replicates were performed. N-number = 26-37. Statistical significance was tested with a paired two-sample Wilcoxon test (n.s., not significant, is given for p-values ≥ 0.05; one star (*) for p-values < 0.05 and ≥ 0.005; two stars (**) is given for values < 0.005 and ≥ 0.0005; three stars (***) is given for values < 0.0005). P-values are shown in Supplementary Table S5. **(D)** Representative images of the rF3H assay for each condition. The area of the LacO locus is highlighted in the mCherry channel where the accumulation of mCherry proteins in the LacO is visible. Scale bar = 5 µm. **(E)** Heatmaps showing ipTMs and pTMs scores of AlphaFold protein-protein interaction predictions between TET proteins and splicing factors without RNA. Higher-ranked predictions shown in Fig. 2 were repeated without the short RNA, to show RNA-specific effects on the prediction score. **(F)** Structure showing AlphaFold 3 prediction model for TET1-CD and DNA (LINE1 5’ UTR region) versus TET1 interaction with RNA (intron-exon transition region).

Next, we investigated whether TET1 proteins affect hm5C levels in J1 mESCs wild-type and different TET1 deletion mutants (Fig. S10A). Previous studies have shown that hm5C levels in embryonic stem cell RNA depend on the presence of TET proteins, decreasing significantly in TET null mice (Fu et al., 2014). Therefore, we quantified hm5C levels in total RNA extracted from wild-type mESCs by slot blot analysis. We performed these experiments by comparing untreated RNA samples with samples boiled at 65 °C, RNA samples treated with RNase A, and plasmid DNA as negative controls. Interestingly, quantification of hm5C levels showed a two-fold increase for samples boiled at 65 °C compared with unboiled samples (Fig. S10B). This suggests that hm5C in RNA is likely found in temperature-sensitive secondary structures. Similar results were published for f5C (Wang et al., 2016), which was shown to favor RNA duplex structures. In addition, we investigated whether different structural domains of TET1 affect the abundance of hm5C in RNA. In addition, we performed slot-blot using total RNA isolated from wild-type mESCs and mESCs expressing different TET1 deletion mutants (Supplementary Table S2). The quantification showed no major changes in hm5C levels in the total RNA of these cell lines. RNA hydroxymethylation levels were not negatively affected by TET1 N-terminal deletions in these cell lines (Fig. S10C), that still share a catalytic domain. In agreement with our previous results, this domain alone is sufficient for RNA binding and m5C oxidation. In addition, all TET1 N-terminal deletion mutants showed a clear colocalization of TET1 with SC35 (Fig. S10D). The studies performed in TET1 deletion mutant cells corroborated AlphaFold structural models, which identified a region in the CRD domain of TET proteins to be involved in their interaction with U2AF. Altogether, this set of experimental approaches verified the interaction between TET1 catalytic domain (TET1-CD) and RNA, demonstrating the role of this protein in RNA metabolism and modifications.

### TET proteins and m5C to hm5C oxidation promote splicing in vitro

Finally, we investigated whether TET proteins and/or their catalytic activity can affect RNA splicing. For this purpose, we used the nonsense-mediated decay (NMD) reporter for alternative splicing in mESCs wild type and knockouts for all three TET proteins or Dnmt. Alternative pre-mRNA splicing is a fundamental regulatory process for most mammalian multi-exon genes to increase proteome diversity. Nonsense-mediated mRNA decay (NMD) is a conserved mRNA surveillance mechanism to mitigate deleterious effects caused by gene mutations or transcriptional errors (reviewed in (Behm-Ansmant et al., 2007). This pathway is initiated by the removal of both the poly(A) tail and the 5′ cap. The latter was shown to be coupled with translational repression (Coller and Parker, 2005). NMD affects mRNAs containing premature translation termination codons (PTCs), which recruit the ATP-dependent RNA helicases Upf1-34. Based on the NMD system and using a transitory transfected NMD reporter vector for EGFP, if alternative splicing takes place, the mRNA encoding EGFP will be translated, and fluorescence intensity in the cells can be measured. If alternative splicing does not occur, the resultant non-sense mRNA will be degraded and EGFP will not be expressed (Fig. 6A). We transfected different cell lines (wild type, TET triple knockout, and DNMT triple knockout) with a vector encoding the NMD reporter and different cherry-tagged TET constructs, including TET1-CDm (catalytically dead mutant), TET1-CD, TET2-CD, and TET3-CD. TET1-CDm harbors a mutation in the Fe(II)-binding motif and maintains its DNA binding ability but is catalytically inactive. We used the catalytic domain of TET proteins based on our previous protein-protein-RNA interaction results (Fig. 2, Fig. 5, and Fig. S10). In addition, this domain is sufficient to oxidize m5C in single-stranded RNA (Fu et al., 2014). After ectopic expression of the vectors, we found an increase in EGFP intensity for TET triple KO cell lines expressing TET1, TET2, and TET3 proteins, including TET1 catalytic dead mutant. Higher EGFP fluorescence relates to higher splicing levels for TET-transfected cell lines compared with controls transfected with mCherry only (Fig. 6B).

**Figure 6.**
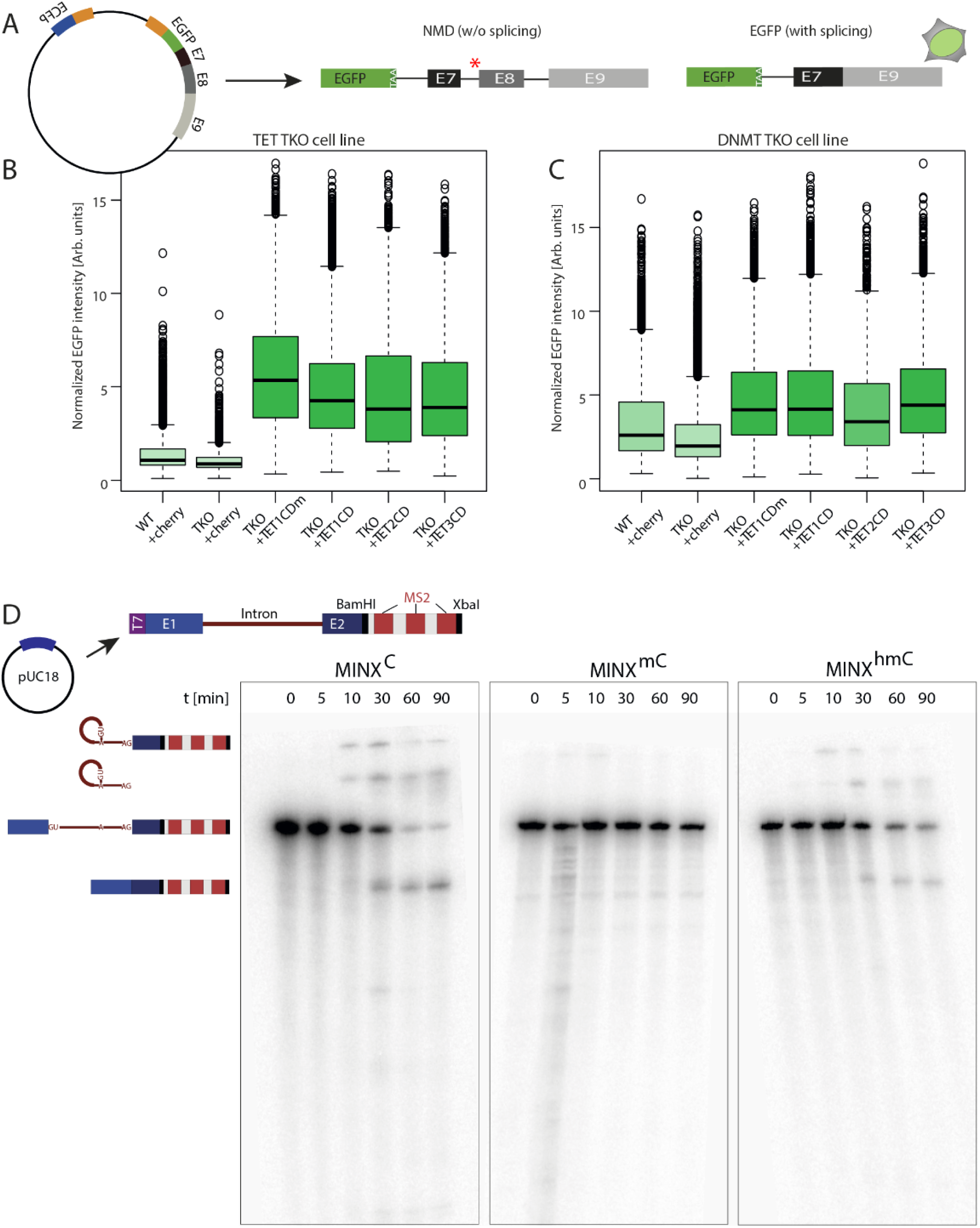
TET proteins and m5C to hm5C oxidation promote splicing. **(A)** Scheme showing the chimeric vector used for the splicing assay based on the NMD system (Non-sense mediated mRNA decay). E7, E8, and E9 correspond to exons 7, 8, and 9 of the HNRNPDL gene (heterogeneous nuclear ribonucleoprotein D-like), represented by black/grey rectangles. Introns between the exons are represented by a black line. The red star indicates premature translation termination codons (PTCs) present in the chimeric “pre-mRNA”. If splicing takes place, the PTC is removed, and translation of the RNA, with GFP at the 5’, occurs. In the absence of splicing, the PTC leads to degradation of the “pre-mRNA”. Stop codon “TAA” at the end of the GFP coding sequence. **(B)** Boxplots showing the normalized EGFP intensity in different mESCs (wild type and TET triple knockouts) transfected with the NMD vector and the different TET constructs. The same experiments were performed in Dnmt triple knockout cell lines, and quantification is shown in **(C)**. **(D)** In vitro splicing assay. On the top is the scheme of the vector used for in vitro splicing. The pre-mRNA encoding adenoviral major late gene sequence exons 1 and 2 interrupted by an intron was created by T7 RNA polymerase run-off transcription. MS2 bacteriophage coat protein binding aptamer (not relevant for this study). For this reaction, the only cytidine sources were either CTP, m5CTP, or hm5CTP. 32p-labeled transcripts were phenol-chloroform purified, and similar RNA amounts were subjected to in vitro splicing reactions with HeLa nuclear extracts (0, 5, 10, 30, 60, 90 min). A diagram representing the different splicing intermediates of this construct is shown on the bottom left. Samples were taken at indicated time points and analyzed on polyacrylamide gels.

We further investigated whether the effect of TET proteins promoting splicing was affected by DNA/RNA 5-methylcytosine levels (5mC/m5C). Both 5mC and m5C can be the substrate of TET proteins, necessary for 5mC/m5C to 5hmC/hm5C conversion. On the other hand, DNMT proteins are responsible for 5mC production, and it has been shown that DNMT2 has double-substrate specificity and adopts a similar catalytic mechanism to methylate RNA (H. Li et al., 2024). Therefore, we used mESCs triple knockouts of DNMT proteins, with residual levels of 5-methylcytosine (Okano et al., 1999; Tsumura et al., 2006). We investigated whether the transitory transfection of TET proteins (as in Fig. 6B) positively affected splicing levels in the absence of 5-methylcytosine. Following the trend of the previous experiment, we also found an increase in the EGFP intensity compared with controls (only mCherry transfected cells) corresponding with an increase in splicing in the presence of TET proteins. In this case, the fold increase was slightly lower than in Fig. 6B using TET triple knockout cell lines (Fig. 6C). This can be explained by higher levels of endogenous TET proteins in these cell lines, already affecting the splicing of the NMD reporter to certain levels, or due to the effect of TET-mediated m5C oxidation favoring splicing in addition to TET presence alone. Mouse ESCs transfected with TET1-CD or TET1-CDm (catalytically dead mutant) showed similar intensity values for EGFP, meaning increased splicing for both (Fig. 6B-C). Therefore, the first option seems more likely. Nonetheless, the levels of alternative splicing still increase for all TET-transfected cells independently of the absence of 5-methylcytosine (in triple DNMT KO cells) or TET1 catalytic activity (TET1-CDmutant). These results show that TET proteins can promote splicing independently of their catalytic activity.

Alternatively, we studied splicing levels using a different splicing reporter system, the spliceable firefly luciferase with one or two introns (human hemoglobin alpha [hHBA] and human hemoglobin beta [hHBB]) under the control of a CMV promoter. Using this system, the relation between the number of introns and splicing performance can be studied in more detail (Fig. S11A). HEK cells were co-transfected with firefly luciferase and mCherry, mCherry-TET1-CD, or mCherry-TET1-CDm. To quantify splicing events, extracts of cells ectopically expressing the desired constructs were mixed with a substrate metabolizable by luciferase. When the mRNA is spliced correctly, the enzymatic activity of luciferase results in a luminescent signal. Then, this luminescence was measured and normalized to the fluorescent signal of mCherry. In these experiments, the luminescence signal was low irrespective of the number of introns for control cells (transfected only with mCherry). However, there was an increase in luminescence signal in cells transfected with both TET1-CD and TET1-CDm, corresponding with an increase in splicing efficiency (Fig. S11B-C). The luminescence signal was much higher when the mRNA comprised a single intron (green bars), compared with the mRNA with two introns (blue bars), indicating that splicing efficiency is almost three-fold higher when only one intron has to be removed. In this case, splicing efficiency is slightly favored by TET1 catalytic activity compared to TET1-CDm. These results suggest that the binding of TET1 alone influences the splicing efficiency of luciferase mRNA, independently of the catalytic activity of TET1. In summary, TET proteins interact with U2AF promoting splicing, an activity that is neither dependent on their catalytic activity nor the substrate (m5C) (Fig. 6 and Fig. S11). However, TET catalytic activity and oxidation of m5C may still play a specific role in favoring splicing efficiency.

To further investigate this, we studied the effect of cytosine modification levels (m5C and hm5C) in RNA splicing. The luciferase splicing reporter showed a slightly higher increase in splicing efficiency for TET1-CD compared to TET1-CDm (Fig. S11B). Therefore, we investigated whether RNA methylation levels and m5C oxidation specifically affect splicing efficiency. For this purpose, we performed in vitro splicing assay using MINX-M3 pre-mRNA reporter construct (Deckert et al., 2006). This vector consists of a pre-mRNA encoding adenoviral major late gene sequences exons 1 and 2 interrupted by an intron and tagged with 2xMS2 bacteriophage coat protein (Fig. 6D). For this purpose, MINX-M3 pre-mRNA was produced with run-off IVT (in vitro transcription) reactions containing either CTP, m5CTP, or hm5CTP, as the only cytidine. 32P-labelled transcripts were subjected to in vitro splicing using HeLa nuclear extracts, a tumor-derived cell line with high levels of TET1 protein (Good et al., 2017). Time course of the splicing reaction was analyzed on polyacrylamide gels (Fig. 6D). Similar splicing kinetics and efficiency were observed for the unmodified and hm5C-containing pre-mRNAs, while no splicing was detected when pre-mRNA containing m5C was used. These results indicate that cytosine methylation affects splicing negatively in vitro, but this can be restored by oxidation of mC to hmC. Thus, RNA modifications like m5C and hm5C also may play a role in regulating splicing efficiency. In a previous publication, DNA base modifications have been shown to affect DNA physical properties and DNA processes, with cytosine methylation stabilizing the DNA helix and increasing its melting temperature. The latter affects DNA helicases and RNA/DNA polymerase speed. In this scenario, the oxidation of methylated cytosines reverses the duplex stability and genome metabolic effects of the unmodified cytosine (Rausch et al., 2021). The latter may be extrapolated to RNA modifications also playing a structural role in RNA metabolism. Together with our finding that hm5C is predominantly found in temperature-sensitive secondary structures, we speculate on the biological role of RNA hydroxymethylation in topologically regulating splicing events. In this way, splicing would be more likely to occur depending on the methylation status of the pre-mRNA, which may form different secondary structures depending on its physical properties. The latter would be related to the variegated deposition of certain cytosine modifications along the pre-mRNA sequence which in turn may rely on TET activity.

## Conclusions and outlook

This study highlights the multifaceted and undisclosed role of TET dioxygenase proteins in RNA metabolism, particularly in regulating RNA splicing in mammalian and Drosophila cells. We demonstrate that TET proteins, beyond their well-established role in DNA demethylation, localize to splicing speckles and interact with key splicing factors such as U2AF1 and U2AF2. These interactions enhance splicing efficiency, independent of TET catalytic activity, as we have shown using different splicing reporter systems and RNA-seq analysis. In this regard, there is previous evidence showing that non-catalytic functions of TET1 are critical for the regulation of gene expression and the silencing of endogenous retroviruses, being an interaction hub for multiple chromatin-modifying complexes (Stolz et al., 2022). However, we also show that TET-mediated oxidation of m5C to hm5C restores splicing efficiency in vitro. This suggests a dual role for TET proteins in splicing: a structural or scaffolding function and a catalytic, RNA modification-dependent function. This dual role may be interconnected, having implications for understanding post-transcriptional and alternative splicing regulation in various biological contexts, including development and disease. Interestingly, it has been shown that a big fraction of RNA splicing regulators are involved in neurogenesis, playing a role in neural development (Fisher and Feng, 2022). In this process, the precise regulation of alternative splicing is crucial (Weyn-Vanhentenryck et al., 2018). TET seems to arise as one of these regulators, as the RNA-seq analysis of splicing events in Drosophila shows. The enrichment of changes on alternative 5’ splice sites for neurodevelopmental genes and the defects in TETnull mutants support this hypothesis.

The distribution of m5C varies among cell types (Amort et al., 2017), and modifications at specific positions on mRNA can have different regulatory roles promoting or inhibiting translation (Hoernes et al., 2016; Li et al., 2017). As we have shown, TET1 association with splicing speckles is RNA dependent, not existing a direct interaction with the splicing factor SC35 (SRSF2). The latter could be interpreted as possible competition of TET and SC35 for binding sites, since this protein has been reported to read m5C in mRNA (Ma et al., 2023). In addition, TET was shown to oxidize m5C in polyadenylated RNAs in Drosophila, favoring mRNA translation. TET and RNA hydroxymethylation were found to be most abundant in the Drosophila brain, with TET-deficient flies suffering from impaired brain development (Delatte et al., 2016). The finding that TET proteins can oxidize 5mC in RNA to promote translation in Drosophila, together with our results in mouse, human, and Drosophila, point to a conserved role of TET proteins in RNA metabolism, affecting splicing efficiency and therefore subsequent translation of transcripts. In this regard, TET proteins arise as a modifier of m5C, in addition to their canonical role in DNA 5mC oxidation. Future research should explore the mechanistic details of TET’s interaction with the spliceosome and the broader impact of RNA hydroxymethylation on RNA structure and function.

In the past, the link between TET1 and the NuRD chromatin remodeling complex was reported (Yildirim et al., 2011), which mediates gene repression to maintain the pluripotent state of ESCs. In addition, we have shown the subnuclear localization of the short isoform of TET1, lacking the zinc finger domain, to heterochromatic regions during the late S phase (Arroyo et al., 2022). These findings suggest that TET1 switches between chromatin-associated and RNA-associated localizations, the latter independent of the cell cycle state. Furthermore, it has been shown that the presence of hm5C in the transcribed gene promotes the annealing of the nascent RNA to the template DNA strand, leading to the formation of an R-loop. The resolution of these R-loops leads to differential expression of a subset of genes involved in stem cell proliferation. In this context, TET activity promotes co-transcriptional R-loop formation as a mechanism of gene expression regulation (Sabino et al., 2022). The latter may be particularly related to our finding that hm5C in RNA seems to be located in secondary structures. Strikingly, TET localization at speckles was found across different cell lines independent of the differentiation state and cell fate. It is also conserved between TET proteins and between mouse, human, and Drosophila cells, the latter without DNA methylation. In this scenario, it is tentative to speculate that the function of TET in splicing reflects its ancestral role, perhaps primary role, in RNA metabolism.

## Material and methods

### Expression constructs

Mammalian expression plasmid encoding mCherry-SC35 (pc3332) was generated by cutting the SC35 coding sequence (CDS) from the vector EYFP-SC35 (pc1202) and cloned into the mCherry-C1 backbone (pc2387) using restriction enzymes BspE1 and HindIII. The vectors pWHE237 (pc3336) and pWHE237mod-WT (pc3337) were used for luciferase reporter gene assays (generated by C. Berenśs laboratory) and published in (Vogel et al., 2018). Both vectors carry the luciferase (luc+) gene. The intron 2 of the human β-globin gene (bgl2 intron) was inserted into the luc+ gene (30 nt downstream of the start codon using the sequence 5′-GTGAGT//CCACAG-3′ as an intron boundary). To generate the splicing reporter vector hnRNP-DL-EGFP_ECFP (pc3351), the firefly luciferase sequence was cut from the vector pWHE200 (pc3335) using BamHI and NotI restriction enzymes, and replaced with EGFP form pc0713, also cut with BspE1 and HindIII. All other expression constructs have been described and published in previous studies and can be found in Supplementary Table S1.

### Cell culture, embryoid body formation, and transfection

Cells were cultured in DMEM (Cat.No.: 41965039, Gibco, Massachusetts, USA) supplemented with fetal bovine serum (FBS) 10% (Capricorn Scientific, Cat.No.: FBS-22A), 2 mM L-glutamate (VWR, Cat.No.: 56-85-9), 0.1% gentamicin (Sigma-Aldrich, Cat.No.: 1405-41-0), sodium pyruvate 110 mg/L (Sigma-Aldrich, Cat.No.: 113-24-6) at 37°C and 5% CO_2_. Mouse embryonic fibroblasts (MEF) were cultured in DMEM containing 15% FBS (Ludwig et al., 2017). Furthermore, the ESC medium for J1 mESCs was supplemented with 16% fetal bovine serum (FBS) (Capricorn Scientific, Cat.No.: FBS-22A), 1x nonessential amino acids (Cat. No.: M7145, Sigma-Aldrich, St Louis, MO, USA), 1x penicillin/streptomycin instead of gentamicin (Cat. No.: P4333, Sigma-Aldrich, St Louis, MO, USA), 2 mM L-glutamine (VWR, Cat.No.: 56-85-9), 0.1 mM β-mercaptoethanol (Cat. No.: 4227, Carl Roth, Karlsruhe, Germany), 1000 U/ ml LIF (Cat. No.: ESG1107, Merck, Kenilworth, NJ, USA), 1 μM of MEK inhibitor PD0325901 and 3 μM of GSK3 inhibitor CHIR99021 (2 inhibitors (2i), Axon Medchem BV, Reston, VA, USA). Stem cells were maintained in a naïve pluripotent ground state by culturing them under feeder-free conditions on gelatin-coated culture dishes (0.2% gelatin in ddH_2_O, Sigma-Aldrich Chemie GmbH, Steinheim, Germany, Cat.No.: G2500) in the above-described stem cell medium. All cells were frozen in freezing medium (DMEM supplemented with 20% FBS, 50 μg/ml gentamicin, and 2 mM L-glutamate, 10% DMSO), and regularly tested for mycoplasma to ensure they were contamination-free. Depending on the experiment, cells were split every two to three days except for ESCs, which were split every two days on gelatinized culture dishes. For in vitro embryoid body (EB) formation J1 ESCs were trypsinized and approximately 37,500 cells/mL were transferred to a fresh tube containing differentiation medium (ESC medium without LIF and 2 inhibitors). Drops of the cell suspension were spotted on the lid of a dish containing 1x PBS (phosphate-buffered saline: 137 mM NaCl, 2.7 mM KCl, 10 mM Na2HPO4, 1.8 mM KH2PO4, in ddH2O, pH ∼ 6.8) in the bottom. After three days EBs were transferred to a new dish containing a differentiation medium and incubated for five days. 24 hours before staining, EBs were trypsinized and seeded on gelatinized coverslips. All cell lines, characteristics, and references are listed in Supplementary Table S2.

HEK-293 cells were transfected using polyethyleneimine (PEI) (pH 7.0, Cat.No.: 40827-7, Sigma-Aldrich Chemie GmbH, Steinheim, Germany) as previously described (Agarwal et al., 2011). For 10 cm diameter dishes 90 µL PEI and 30 µg plasmid DNA were added to 900 µL DMEM without supplements and mixed by vortexing. For 6-well plates, 12 µL PEI and 4 µg plasmid DNA were added to 200 µL DMEM. The mixtures were combined, vortexed for 80 seconds, and incubated at room temperature for at least 30 minutes. Then, the mixture was added dropwise to the cells growing to a confluency of 80% before transfection. For transfection by electroporation, MTF and BHK cells were transfected using an AMAXA Nucleofector® system II (Lonza, S/N: 10700731) using a self-made buffer (5 mM KCl (Sigma-Aldrich Cat.No.: 7447-40-7), 15 mM MgCl2 (Sigma-Aldrich Cat.No.: 7786-30-3), 120 mM Na2HPO4/NaH2PO4 (Sigma-Aldrich Cat.No.: 7558-79-4) pH 7.2, 50 mM Mannitol (Caesar & Loretz, Cat.No.: 69-65-8)) (Becker et al., 2016a) with default programs for each cell line. Cells growing at 60-80% confluency were trypsinized and centrifuged (6.5 min; 1,400 rpm) the day before the experiment. Afterward, cells were resuspended in 100 µL of nucleofection buffer containing 5 µg of the plasmid DNA. Subsequently, cells were seeded on gelatinized coverslips or glass-bottom p35 plates for live-cell experiments (rF3H). Mouse embryonic stem cells were transfected with the Neon electroporation system (ThermoFisher Scientific) according to the manufacturer’s instructions.

### Mass spectrometry sample preparation, LC-MS/MS, and data analysis

GFP-tagged TET proteins were expressed in HEK293T cells. Cell lysis and immunoprecipitation with the GFP-Trap (ChromoTek GmbH, Martinsried, Germany) were performed as described previously (Müller et al., 2014). After co-immunoprecipitation, protein samples were digested on beads with trypsin according to standard protocols. Samples on beads were rinsed two times with wash buffer (20 mM Tris-HCl (pH 7.5), 300 mM NaCl, and 0.5 mM EDTA) and two times with immunoprecipitation buffer (20 mM Tris-HCl (pH 7.5), 150 mM NaCl, and 0.5 mM EDTA). The protocol described in (Bauer et al., 2015) was applied as follows. 100 μl of denaturation buffer containing 6 M guanidine hydrochloride, 10 mM tris (2-carboxyethyl)phosphine, and 40 mM chloroacetamide in 100 m Tris (pH 8.5) was added to the beads and heated at 70 °C for 5 minutes. The samples were then subjected to sonication in a Diagenode Bioruptor Plus system (UCD-300-TO) at maximum power settings for 10 cycles comprising 30-s pulse and 30-s pause. Following sonication, the samples were diluted 1:10 with a digestion buffer (25 mM Tris (pH 8.5) containing 10% acetonitrile) and mixed by vortexing before enzyme digestion. Each sample was digested with 1 μg of endoproteinase Lys-C (Wako Chemicals, Neuss, Germany) for 4 h with subsequent digestion using 1 μg of trypsin (Promega, Madison, WI) under gentle rotation at 37 °C. After digestion, the samples were placed in a SpeedVac concentrator for 10 minutes to remove acetonitrile from the sample before StageTip purification using SDB-XC material (Rappsilber et al., 2003). Peptides were then eluted from the StageTip and placed in the SpeedVac concentrator to reduce the sample volume. Peptide mixtures were analyzed by electrospray MS/MS spectrometry, injecting 5 μl of the sample into the column. Experiments were performed with an LTQ Orbitrap XL mass spectrometer (Thermo Scientific, Waltham, MA). Spectra were analyzed with Mascot software (Matrix Science, Boston, MA).

Samples were loaded onto a column (15-cm length and 75-μm inner diameter; New Objective, Woburn, MA) packed with 3-μm ReproSil C18 beads (Dr. Maisch GmbH, Ammerbuch-Entringen, Germany) using an EASY-nLC autosampler (Thermo Scientific) coupled via a nanoelectrospray source to the LTQ Orbitrap XL mass spectrometer. Each sample was analyzed using a 2-h reversed-phase gradient and a top 5 method for data-dependent acquisition. Full scans were acquired in the Orbitrap mass spectrometer after accumulating up to 1 × 10^6^ charges, and MS/MS with the five most abundant precursors was performed using low-energy ion-trap collision-induced dissociation. MS/MS spectra were recorded using the ion trap by radial ejection.

All raw files were analyzed using the MaxQuant computational proteomics platform (version 1.4.1.6) (Cox and Mann, 2008) as described in (Bauer et al., 2015). Peak lists were searched against the UniProt database (https://www.uniprot.org/) with an initial mass deviation of 7 ppm and fragment ion deviation of 0.5 Thomson. Carbamidomethylation was used as a fixed modification. Oxidation of methionine and acetylation of the protein N terminus were used as variable modifications. All unmodified and oxidized methionine- and N-acetylation-containing peptides were used for protein quantification with the label-free quantitation algorithm. MaxQuant output data were further analyzed with Perseus software (version 1.5.0.15) (Cox and Mann, 2008). Significance was tested using a Student’s two-tailed paired t-test. All experiments were performed in biological triplicates.

### Structural modeling with AlphaFold 3 and AlphaFold 2-Multimer

The AlphaFold3 web interface was used to screen protein-protein-RNA interactions (https://alphafoldserver.com/) (Abramson et al., 2024). To rank the predictions, both pTM and ipTM scores were used. The predicted template modeling (pTM) score and the interface-predicted template modeling (ipTM) measure the accuracy of the entire structure (Zhang and Skolnick, 2004; Xu and Zhang, 2010). A pTM score above 0.5 means the overall predicted fold for the complex might be similar to the true structure. ipTM measures the accuracy of the predicted relative positions of the subunits within the complex. Values higher than 0.8 represent confident high-quality predictions, while values below 0.6 suggest likely a failed prediction. PAE and pLDDT were also taken into consideration after a close inspection of the models. PAE plots for full-length protein predictions and pre-existing models in the 3D structures databases (AlphaFoldDB) were used to perform sequence fragmentation for further predictions. Well-predicted domains (pLDDT > 90) were not fragmented and were taken as a unit core. As an RNA sequence in protein-protein-RNA interactions, we selected a short fragment located in the intron-exon limit of the spliceable firefly luciferase (5’-CCCACAGCCGGCG-3’) (splicing reporter system, pc3336). Protein fragments were designed using available monomeric structural models from AlphaFold as provided by the AlphaFold database (Varadi et al., 2024). To model TET1-DNA interaction, we selected a short DNA sequence in the LINE 1 5’ UTR (5’-GCGCA CCTTC CCTGT AAGAG AGCTT GCCAG CAGAG AGTGC TCTGA-3’) that is hydroxymethylated by TET1s (Arroyo et al., 2022). To model TET1-RNA interaction, we used the following fragment of the splicing reporter pc3336: 5’-AAAACAUAAAGAAAGGCGUGAGUCUAUGGGA-3’. A local installation of AlphaFold-Multimer 2.3.2 was used to perform selected structural modeling with AMBER relaxation (Evans et al., 2021). Five predictions were generated per model, and the prediction with the highest model confidence was used for verification of AlphaFold 3 predictions. All protein sequences were extracted from UniProt (UniProt Consortium, 2023). Data analysis and plotting were done with Python, using pandas and numpy packages. Protein structure images were generated with ChimeraX (Pettersen et al., 2021; Schrodinger, n.d.). All the structural models, generated “.pbd” files, and associated data, including PAE values, “.json”, and scores, are deposited and available at the repository TUdatalib (https://tudatalib.ulb.tu-darmstadt.de/handle/tudatalib/4453).

### Immunostaining of mammalian cells

Immunostainings were performed as described in (Cardoso et al., 1997). For this purpose, cells were seeded on gelatine-coated glass coverslips. Afterward, cells were washed with 1x PBS and fixed in 3.7% formaldehyde (Sigma-Aldrich Chemie GmbH, Steinheim, Germany, Cat.No.: F8775) in 1x PBS for 10 minutes. After fixation and three washing steps with PBS-T (1x PBS, 0.01% Tween-20), cells were permeabilized with 0.5% Triton X-100 in 1x PBS for 20 minutes. Then, cells were washed with PBS and blocked in 0.02% fish skin gelatine or 1% BSA in PBS for 30 minutes to one hour. After blocking, the primary antibodies diluted in blocking solution were incubated for 70 minutes, followed by three washing steps with 0.01% Tween20 in PBS (10 minutes each). Incubation with the secondary antibodies diluted in 1% BSA in PBS was performed for one hour, followed by three washing steps with 0.01% Tween in PBS for 10 minutes each. Finally, DNA was counterstained with DAPI (4,6-diamidino-2-phenylindole, 10 g/ml, Cat. No.: D27802, Sigma-Aldrich Chemie GmbH, Steinheim, Germany) for 10 minutes and samples were mounted in Mowiol 4-88 (Cat. No.: 81381, Sigma-Aldrich Chemie GmbH, Steinheim, Germany) containing 2.5% DABCO (1,4-diazabicyclo[2.2.2]octane, Cat. No.: D27802, Sigma-Aldrich Chemie GmbH, Steinheim, Germany). For immunostaining of mouse tissues, CrK(SO_4_)_2_ coated slides with paraffin-embedded mouse tissue sections were incubated for three hours at 65°C. To remove paraffin 100% xylol (3x 5 minutes) and a series of decreasing ethanol concentrations (100%, 96%, 90%, 80%, 70%; each for 5 minutes) was used. For antigen retrieval, slides were autoclaved (100°C for 30 minutes, end temperature 80°C) in 1 mM Tris-EDTA (pH 8.5) with subsequent incubation in 0.05% Tween in PBS two times for 15 minutes and briefly in PBS. Paraffin-embedded tissues were blocked with 1% BSA in PBS for 30 minutes. Primary antibodies were incubated overnight at 4°C. Secondary antibodies were incubated in 1% BSA for one hour. After antibody incubation, samples were washed with 0.01% Tween in PBS (0, 5, 15 minutes). DNA was counterstained using DAPI, with a subsequent washing step using PBS and ddH_2_O. Slides were mounted in Mowiol. Cryosections on microscopy slides (SuperFrost Ultra Plus, Roth, Germany) were dried up for 30 minutes and transferred to 10 mM sodium citrate buffer (pH 6.5) for 3 minutes. For antigen retrieval, slides were heated up to 80°C (water bath) in the same solution. Then, incubated in 0.5% Triton X-100 in PBS for one hour. Primary and secondary antibodies in a blocking solution containing 20 µg/mL DAPI, were incubated in humid dark chambers (Arroyo et al., 2023) overnight. Wash steps were performed using 0.01% Triton X-100 in PBS three times for 30 minutes at 37°C. Slides were mounted in Vectashield. For RNase treatments followed by extraction and immunofluorescence, MTF cells were seeded on glass coverslips. 24 hours later cells were washed once with Triton 0.01% in PBS and incubated for 7 minutes with RNaseA 10 mg/ml in PBS. After RNaseA treatment, cells were fixed with paraformaldehyde 3.7% in PBS and immunostained with antibodies against TET1 and SC35. Subsequent steps were performed as described before. As treatment control, cells subjected to the same treatments (extraction and RNaseA incubation) were stained with propidium iodide (Cat. No.: P4170, Sigma-Aldrich) 1:200 in 1x PBS, and the respective signal was quantified by high-content microscopy (n>1000). All primary and secondary antibody details, as well as dilutions used, are available in Supplementary Table S3.

### Immunostaining of Drosophila salivary glands cells

For the analysis of colocalization SC-35 and RNA Pol II in the salivary gland of the larvae of Drosophila, tissues were dissected in PBS, fixed in 4% paraformaldehyde in PBS for 20 minutes at RT, washed in PBST (PBS with BSA and 0.3% Triton-× 100). Antibody in-situ stainings were done as described previously (Haussmann et al., 2008) using rabbit anti-HA (1:1000; Sigma), mouse anti-SC35 (Sigma-Aldrich S4045 ascites, 1:1000), rat anti-RNA Pol II (anti-Ser 2, clone 3E10, 04-1571 Merck-Millipore, 1:1000) and visualized with Alexa Fluor 488 (1:250; Molecular Probes), Alexa Fluor 546 (1:250; Molecular Probes) or Alexa Fluor 647 (1:250; Molecular Probes). For imaging, tissues were mounted in Vectashield (Vector Labs) and visualized with confocal microscopy using a Leica TCS SP8. Images were processed using Fiji (https://Fiji.nih.gov/ij/).

Recombineering was used to clone the genomic part of the core TET gene into pUC 3GLA UAS from a genomic BAC clone as described (Dix et al., 2024; Haussmann et al., 2019) by adding a HA Tev FLAG c-terminal tag. The N-terminal part for each isoform was then added from PCR-amplified cDNA. For the catalytically dead mutant short isoform, the conserved HxD iron binding motif required for the catalytic activity of TET/AlkB dioxygenase family enzymes was mutated to AxA (Hu et al., 2013). Transgenic lines were generated by phiC31-mediated insertion into the attP2 landing site. For immunohistochemistry, the GFP marker was removed by Cre recombinase.

### RNA-seq analysis

The analysis of alternative splicing and differential gene expression in Drosophila TETnull mutants was done as previously described (Boulet et al., 2023; Haussmann et al., 2022) using published RNAseq data (Boulet et al., 2023; Haussmann et al., 2022). RNA-seq data sets can be found in GEO under the accession number GSE116212. GO analysis was done with STRING using the rank list enrichment analysis.

### Microscopy and image analysis

Confocal images were acquired using a Nikon Eclipse Ti microscope controlled by the software Volocity 6.3 and equipped with an UltraView VoX spinning disk system (Perkin Elmer; Waltham, MA) and a Hamamatsu C9100-50 cooled EMCCD14 camera. As an objective, a Nikon CFI Plan Apo VC 60x oil immersion (60/1.44 NA) and lasers with excitation lines 405, 488, 561, and 640 nm were used. Z-stacks were acquired with a voxel size of XY=120 nm and a Z-step length of 0.3 µm. In addition, Leica TCS SPE-II and SP5 systems mounted on a DMi8 stand and controlled by the Leica LAS X software were used for image acquisition. As objectives, a Leica ACS APO 20x/0.60 NA corrected for oil immersion and Leica ACS APO 63x/1.30 NA Oil CS 0.17/E,0.16 were used. Laser with 405, 488, 561, and 635 nm were used. Images of mouse tissues were acquired using a Leica TCS SP5 confocal microscope (Milton Keynes, UK) equipped with Plan Apo 63x/1.40 NA oil immersion objectives and lasers with excitation lines 405, 488, and 633 nm. For computational image analysis, images were analyzed with Fiji (http://fiji.sc/). A Gaussian Blur filter with a sigma radius of 1.0 was applied to all images. For a more detailed colocalization analysis, the Python-based image analysis platform Priithon (http://code.google.com/p/priithon/) was used to calculate the H-coefficient (Herce et al., 2013). Accumulation and rF3H assay image analysis was performed as described in (Arroyo et al., 2022). To image and quantify splicing reporter assays, the Operetta high-content screening system (Perkin Elmer, UK) was used in wide-field mode, equipped with a Xenon fiber optic light source and a 20×/0.45 NA long working distance or a 40×/0.95 NA objective. For excitation and emission, the following filter combinations were used, 360–400 nm and 410–480 nm for DAPI, 460–490 nm and 500–550 nm for Alexa-488 as well as 560–580 nm and 590–640 nm for Alexa-594. Fluorescence intensity levels were quantified with the Harmony software (Version 3.5.1, PerkinElmer, UK).

### RNA fluorescent three-hybrid assay (rF3H)

We performed the RNA fluorescent three-hybrid assay to address the interaction between TET1 proteins and RNA, specifically an mRNA mimic (Duan et al., 2021). To this end, BHK cells were transfected with plasmids encoding an RNA trap fused to EGFP and the LacI protein for targeting the LacO present in this cell line (pc4555), an mRNA mimic (ms2-PABCP1-mCherry, pc4573), and mCherry-TET1-CD (pc2547). As a positive control of the assay, cells were transfected with the RNA trap, the mRNA mimic, and PABPC1-mCherry which was shown to interact with the mRNA mimic (Duan et al., 2021). As a negative control, only the mRNA trap plus PABPC1-mCherry were transfected (Fig. 5B). Cells were imaged live 8–12 h post-transfection and confocal Z-stacks (voxel size, 0.12 × 0.12 × 0.5 µm) were acquired using the confocal microscope Leica TCS SP5 II. Z-stacks were analyzed and mounted using Fiji (https://Fiji.nih.gov/ij/). Cells showing GFP and mCherry signals were selected. The LacO was segmented in each nucleus for image analysis using the GFP signal and generating an ROI (region of interest). Then, sum intensities for mCherry proteins were measured in this ROI. A second ROI of the same shape and size was used to measure the sum intensity of mCherry proteins in a region outside the LacO. The relative accumulation at the LacO was calculated as the ratio between mCherry sum intensity in the LacO and mCherry sum intensity outside the LacO.

### Luciferase and NMD splicing reporter assay

For luciferase splicing reporter assay, HEK cells were transfected with PEI using 2 µg of mCherry-tagged TET1 constructs (TET1-CD/TET1-CDmut) or mCherry alone, in combination with 2 µg of plasmid DNA encoding firefly luciferase constructs with either one (pc3336) or two introns (pc3337). 24 hours post-transfection, the cell culture medium was removed and the cells were washed with 1x PBS. 200 µL of Cell Lysis Reagent (Luciferase Assay System, Promega) was added and incubated for 10 minutes at room temperature. Cells were scraped and transferred to a new 1.5 mL tube. 50 µL of the cell extract was mixed with 100 µL of Luciferase Assay Reagent (Luciferase Assay System, Promega) in a 96-well plate and luminescence was measured immediately for 20 seconds per well using a microplate reader (Infinite® 200 PRO series, TECAN). Afterward, the fluorescence signal intensity of mCherry was measured for the whole plate and used for normalization of the luminescence signal. For NMD reporter assay with EGFP, J1 mESC wild type and TET or Dnmt triple knockout cell lines were transfected with N-terminal mCherry tagged catalytic domains of TET1, TET2, and TET3. mCherry alone was used as control. In this assay, the mRNA is translated when the exon8 at RNP-DL is spliced out. If no splicing occurs, the mRNA is degraded and no fluorescence signal is detected. The intensity of the EGFP signal was quantified by microscopy and image analysis using the Operetta high-content microscopy system (Perkin Elmer, UK) in wide-field mode, equipped with a Xenon fiber optic light source and a 20x/0.45 NA long working distance objective. For excitation and emission, the following filter combinations were used, 360-400 nm and 410-480 nm for DAPI, 460-490 nm for GFP, and 560–580 nm for TexasRed (mCherry). Fluorescence intensity levels were quantified with the Harmony software (Version 3.5.1, PerkinElmer, UK)

### In vitro splicing assay

In vitro splicing experiments were performed with MINX-M3 pre-mRNA (Deckert et al., 2006; Zhang et al., 2017; Zillmann et al., 1988). DNA templates for run-off IVT (in vitro transcription) reactions were generated via digesting the plasmid with XbaI restriction enzyme. MINX-M3 pre-mRNAs with different methylation status of cytosines were in vitro transcribed in 50µl reactions containing T7 transcription buffer (5X stock: 600 mM HEPES-KOH pH 7.5, 160 mM MgCl2, 10 mM spermidine, 200 mM DTT (Dithiothreitol)); 7.5 mM ATP; 7.5 mM CTP, mCTP, or hmCTP; 1.5 mM GTP; 1.5 mM UTP; 5 mM 3mGpppG cap analog; 5µl 32P-UTP (3000 Ci/nmol, 10 μCi/μL); 50 ng/μL template DNA; 6 μL T7 polymerase, 2 μL RNasin and 0.5 μL YIPP IVTs were performed for 2h and 37 °C, and RNA was subsequently purified via 4% denaturing PAGE and extracted with phenol-chloroform. In vitro splicing reactions with IVT pre-mRNAs were performed for 0-90 minutes at 30 °C in the presence of 40% HeLa nuclear extract, 3mM MgCl_2_, 65mM KCL, 20mM HEPES-KOH pH 7.9, 2mM ATP and 20mM creatine phosphate). RNA was extracted from splicing reactions with protein K digestion followed by phenol-chloroform extraction and ethanol precipitation. Isolated RNA was then analyzed on 14% denaturing PAGE and visualized with autoradiography.

### Co-immunoprecipitation and Western blotting

Co-immunoprecipitations were performed as described before (Becker et al., 2016b). To analyze the protein-protein interaction between TET and splicing factors, transfected HEK cells were harvested by trypsinization and centrifuged for 10 minutes at 2,000 rpm and 4°C. The pellet was resuspended in 200 µL ice-cold lysis buffer containing 150 mM NaCl, 20 mM Tris HCl (pH 8), 1.5 mM MgCl2, 0.2 mM EDTA, 0.4% NP-40, and protease inhibitors 1 mM AEBSF (4-(2-Aminoethyl) benzyl sulfonyl fluoride hydrochloride, Cat. No.: A1421.0100, VWR, Radnor, PA, USA), 1 mM E64 (Cat. No.: E3132, Sigma-Aldrich, St Louis, MO, USA), 1 nM Pepstatin A (Cat. No.: 77170, Sigma-Aldrich, St Louis, MO, USA), PMSF (10 µM, Sigma-Aldrich, St. Louis, MO, USA/Solarbio; catalog #P8340) and AEBSF (1 mM, AppliChem, Darmstadt, Germany). Cells were homogenized with a syringe (21 G needles, 20 strokes) and incubated on ice for 30 minutes with repeated vortexing in between. Lysates were then cleared by centrifugation for 15 minutes at 13000 xg and 4 °C. 15% of the lysate was used as input and the rest was incubated with GFP-binder beads produced as described before (Rothbauer et al., 2008) on a rotator at 4 °C for 90 minutes. A buffer containing 3% PMSF and 1 mM Ribonucleoside Vanadyl Complex RNase Inhibitor (NEB) was added to the beads. Afterward, the beads were washed three times with 500 µL washing buffer (150 mM NaCl, 20 mM Tris HCl (pH 8), 1.5 mM MgCl2, 0.2 mM EDTA). Input and bound fractions were boiled at 95 °C in 4x SDS loading buffer (200 mM Tris/HCl pH 6.8, 400 mM DTT, 8% SDS, 0.4% bromophenol blue and 40% glycerol), separated on 8% SDS-PA (sodium dodecyl sulfate–polyacrylamide) gels. SDS-PAGE and Western blotting were performed as described in (Mortusewicz et al., 2006). Co-immunoprecipitation samples were transferred onto a nitrocellulose membrane (GE Healthcare, München, Germany) in a semi-dry blotting chamber (Bio-Rad Laboratories) for 50 minutes at 25 V. The transfer was verified by Ponceau S staining. Blocking of membranes was performed for one hour in 3% low-fat milk in 1x PBS, followed by incubation with primary antibodies diluted in blocking buffer overnight at 4 °C. The membrane was incubated with the respective secondary antibodies after washing 3 times (10 minutes each) using 1x PBS supplemented with 0.01% Tween-20. Horseradish peroxidase (HRP) conjugated secondary antibodies were used. Characteristics of all primary and secondary antibodies, as well as dilutions used, are described in Supplementary Table S3. Visualization of immunoreactive bands was achieved by Pierce™ ECL Western Blotting Substrate (Cat. No.: 32209, ThermoFisher Scientific, Waltham, MA, USA). For imaging, the Amersham AI600 Imager with a CCD camera was used (GE Healthcare, Chicago, II, USA). Cutouts of the membranes were made for a better composition of the figures. Uncropped and unprocessed scans for all the blots are available, including replicates in supplementary figures, and are provided with the data sets uploaded to TuDatalib (https://tudatalib.ulb.tu-darmstadt.de/handle/tudatalib/4453).

### RNA isolation and slot blotting

To investigate the hm5C amount in J1 ESCs, total RNA was isolated using a Qiagen RNeasy® Kit. Isolation was mainly performed following the manufacturer’s instructions with the following changes. Cell pellets were vortexed with 350 µL RLT buffer and passed 10 times through a blunt 20G needle fitted on an RNase-free syringe. 350 µL of 70% ethanol (in DEPC water) was added and mixed by pipetting. 700 µL were transferred to an RNeasy® spin column and centrifuged for 15 seconds at 10,000 rpm. At the end of the protocol, RNA was eluted twice with 30 µL RNase-free water. After RNA isolation DNA was digested using rDNase from Nucleospin® Triprep Kit (Macherey Nagel). For RNA clean-up samples were filled up to 100 µL and mixed with 350 µL RLT buffer containing 2-mercaptoethanol (1:100). 250 µL of 100% ethanol was added and mixed by pipetting. 700 µL were transferred to an RNeasy® column (Qiagen) and centrifuged for 15 seconds at 10,000 rpm. Afterwards, 500 µL RPE buffer was added and samples were centrifuged again. After discarding the flowthrough 500 µL 80% ethanol (in DEPC water) was added and samples were centrifuged for 2 minutes at 10,000 rpm. Columns were placed into fresh tubes and RNA was eluted with 20 µL RNase-free water by centrifugation for 1 minute at 13,200 rpm. To assess the concentration and purity of RNA, the ratio of absorbance at 260 and 280 nm was measured on a TECAN infinite M200 plate reader (Tecan Group Ltd., Maennedorf, Switzerland).

For slot blot analysis of hm5C in total RNA, isolated samples were blotted on nylon membranes pre-equilibrated with 20x SSC. For controls, plasmid DNA was digested with HindIII and EcoRI (20 µL total volume, NEB buffer 2.1) for 1 hour at 37°C, and an RNA sample was digested with RNase A (20 µL/mL) for 30 minutes at 37°C. Respective RNA amounts were applied in duplicates to the slots of a blotter (Schleicher & Schuell) and pulled through by vacuum. After air drying, the membrane was UV-crosslinked with 120,000 µJ/cm2 two times for 1 minute. To calculate the amount of membrane-bound RNA, half of the samples were stained with 0.02% methylene blue in 0.3 M sodium acetate. The other half was blocked with 3% milk in PBS for 30 minutes. The 5hmC antibody was incubated overnight at 4°C, and the secondary antibody for one hour at room temperature. Following antibody incubations the membrane was washed with 0.1% Tween in PBS (3× 10 minutes). To detect the chemiluminescent signal, a Luminol Solution (Thermo Scientific, Pierce ECL Plus Western Blotting Substrate) was applied to the membrane and images were acquired with an Amersham Imager 600 (Amersham).

### Statistics and Reproducibility

Data visualization and statistical analysis were performed using RStudio (versions V1.2.5033 and V2023.03.1-446, https://rstudio.com/) and MicrosoftⓇ ExcelⓇ for Mac 2011 (Version 14.7.7) unless stated otherwise. Barplots show the average value of the distribution and the whiskers represent the standard deviation with a 95% confidence interval. In all figures showing boxplots, the box represents 50% of the data, starting in the first quartile (25%) and ending in the third (75%). The line inside represents the median. The whiskers represent the upper and lower quartiles. In most of the plots, outliers are excluded and defined as 1.5 times the interquartile range. For the statistics, an independent two-group comparison was made for some conditions with Wilcoxon-Mann-Whitney or One-Way ANOVA tests. Related to this, n.s., not significant, is given for p-values > or equal to 0.05; one star (*) is given for p-values < 0.05 and > or equal to 0.005; two stars (**) is given for values < 0.005 and > or equal to 0.0005; three stars (***) is given for values < 0.0005; between the top of two boxes subjected to comparison. Statistical values (number (#) of cells (N), mean, median, standard deviation (SD), standard error of the mean (SEM), 95% confidence interval (CI), and p-values are summarized in Supplementary Table S5. The cells analyzed showed the reported behavior of the representative images selected.

## Supporting information

Hastert et al. 2025 Supplementary information

Supplementary Data S1 - Mass spectrometry

Supplementary Data S2 - AlphaFold3 scores

Supplementary Data S3 - RNA-seq differential gene expression analysis

Supplementary Data S4 - RNA-seq splicing events analysis

## Data and Code Availability Statement

Renewable biological materials will be made available upon request from the corresponding author Maria Arroyo and M. Cristina Cardoso. All our data sets, including unprocessed images and source datasets have been deposited and are available at TuDatalib (https://tudatalib.ulb.tu-darmstadt.de/handle/tudatalib/4453). Published RNA-seq data sets used in this study can be found in GEO under the accession number GSE116212.

## Acknowledgments

We thank the bioinformatics core facility of the Institute of Molecular Biology (IMB) for providing support in AlphaFold 2 modeling. We thank Prof. Dr. Reinhard Lührmann for his advice and help with the in vitro splicing assay. We also thank Prof. Dr. Florian Heyd, Prof. Dr. Beatrix Suess, and Dr. Marc Vogel for kindly providing relevant materials for this study. We thank BacPac for BAC clones, L. Waltzer for TET mutants, the Bloomington stock center for flies, and FlyORF for injections.

## Author contributions

F.D.H., J.W., C.B., M.A., A.Z., D.N.D.S., T.C.D, and R.A. performed experiments. F.D.H., J.W., C.B., M.A., A.Z., D.N.D.S., T.C.D, and R.A. analyzed data. F.D.H. developed data analysis pipelines. M.A. performed structural modeling with AF and data analysis. S.B., H.L., and M.S. provided tools and advice. F.D.H., M.A., and M.C.C. conceived and developed the project. M.A. generated the final figures and wrote the manuscript. All authors agreed on the manuscript and contributed to the editing of the manuscript.

