## Supplementary material for "TET dioxygenases localize at splicing speckles and promote RNA splicing": Hastert et al. 2025 Supplementary information

#### Supplementary figures and legends

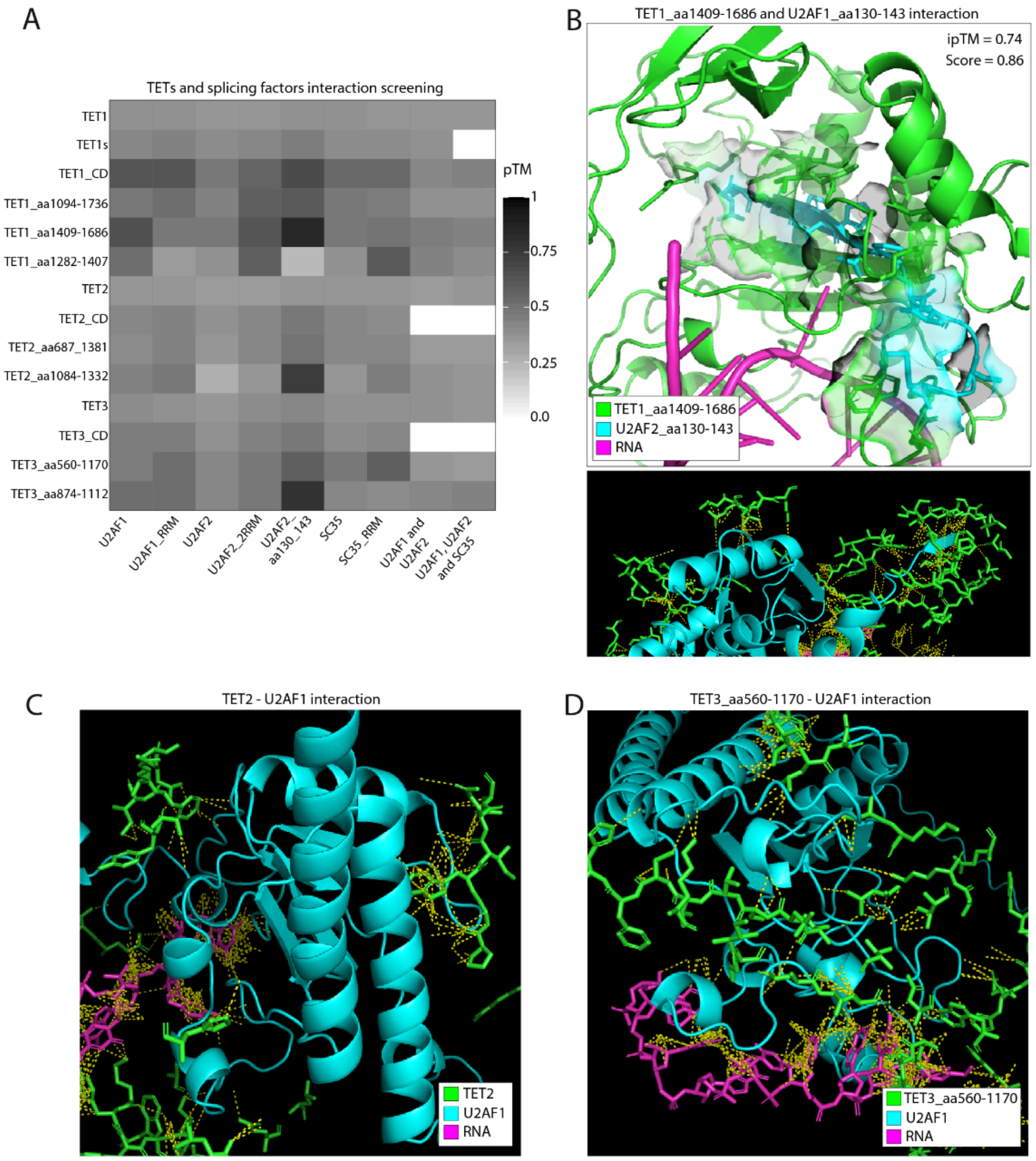

**Figure S1. Screening of TET protein interaction with splicing factors using AlphaFold 3. (A)** Heatmap showing the pTM score (predicted Template Modeling score) obtained for AF structure models for all the interactions tested between TET proteins and the splicing factors U2AF1, U2AF2, and SC35, together with RNA. Labels on the x and y axes indicate the paired protein fragments for structural modeling. White tiles indicate pairs that were not subjected to structural modeling. Magnification of the highest-scored structural models obtained for **(B)** TET1-U2AF2-RNA, **(C)** TET2-U2AF1-RNA, and **(D)** TET3-U2AF predictions. TET residues within a maximum distance of 5 angstroms to U2AF proteins are shown. All types of contacts between chains in a distance of 4 angstroms are shown as yellow dashed lines.

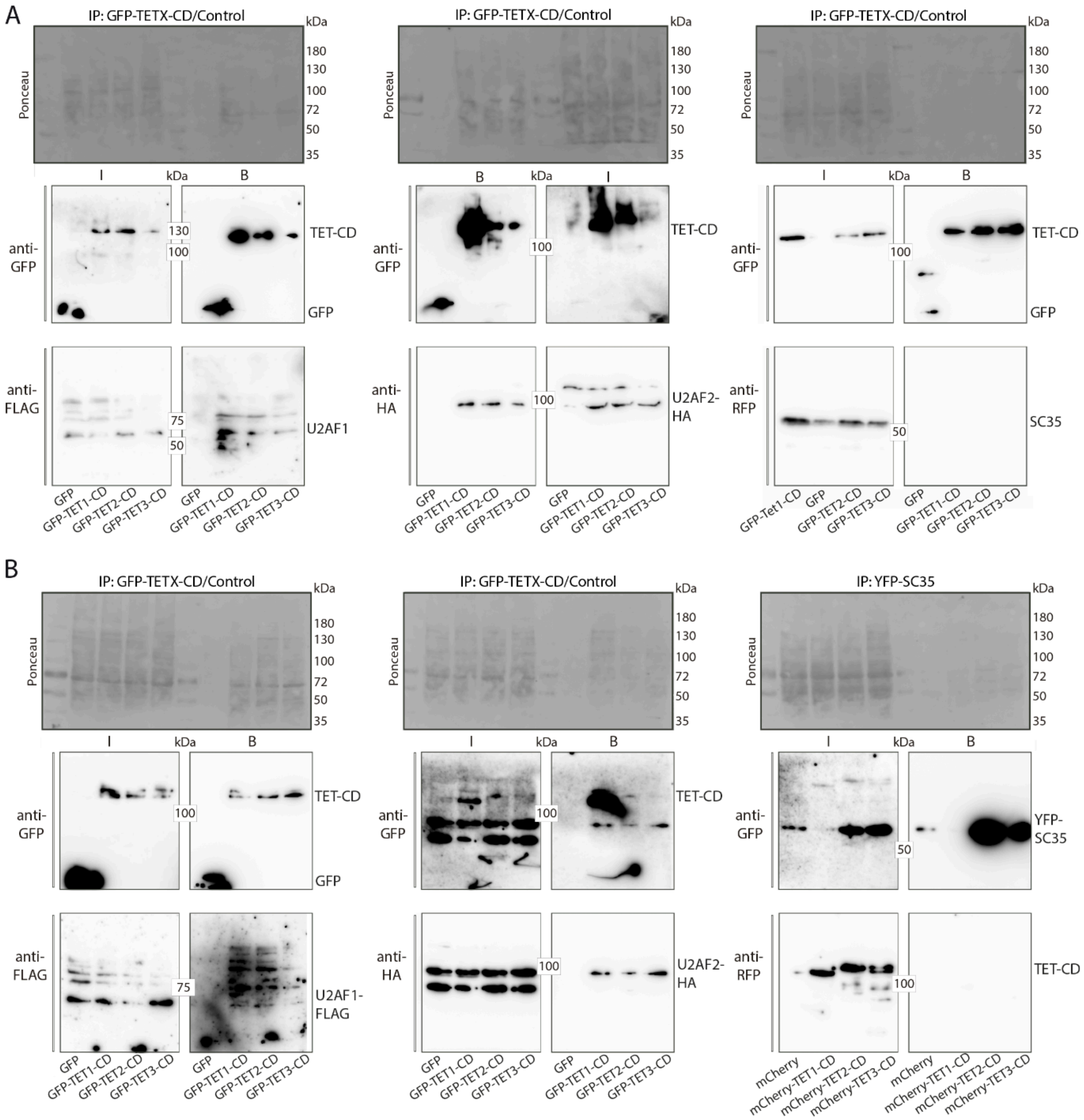

**Figure S2. The catalytic domains of TET1, TET2, and TET3 interact with the splicing factors U2AF1 and U2AF2 but not with SC35. (A)** Co-immunoprecipitation experiments: HEK cells were transfected with EGFP or EGFP-tagged TET1, TET2, or TET2 catalytic domain, together with the splicing factors U2AF1-FLAG, U2AF2-HA, or mCherry-SC35. Cell extracts were analyzed by immunoprecipitation with immobilized GFP-binding nanobodies, followed by detection with antibodies against GFP or FLAG/HA/mCherry. Input "I" and GFP-binding fraction "B" are shown. **(B)** Co-immunoprecipitation experiments were performed as described in (A) for U2AF1-FLAG and U2AF2-HA. For SC35, HEK cells were transfected with YFP-SC35 and mCherry-tagged TET1, TET2, or TET3 catalytic domains. YFP-SC35 was immunoprecipitated with an immobilized GFP-binding nanobody, followed by detection with antibodies against GFP/RFP.

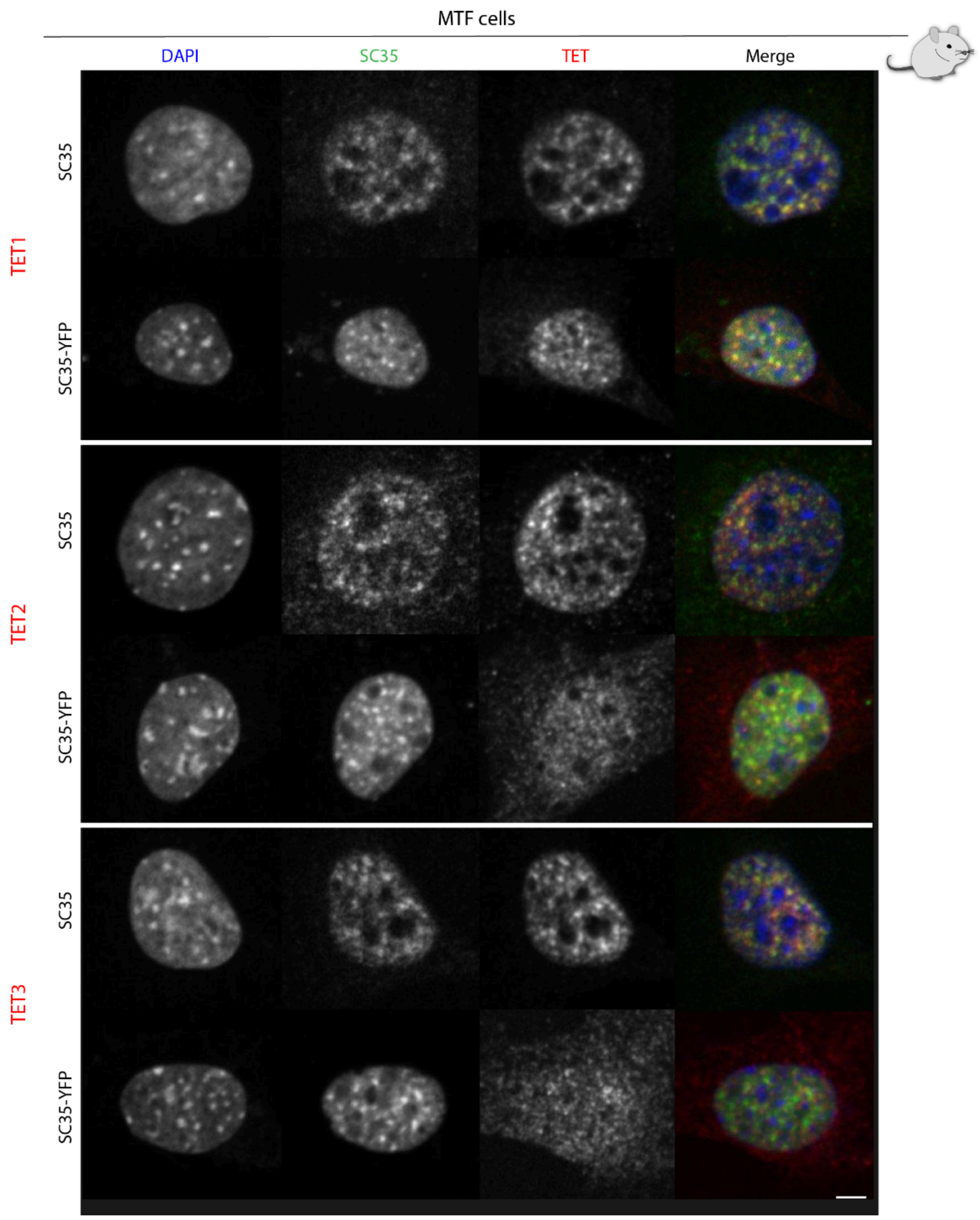

**Figure S3. TET1, TET2, and TET3 localization at splicing speckles visualized with SC35 in mouse cells.** MTF cells (mouse tail fibroblast) were immunostained for endogenous TET proteins and SC35 and imaged by confocal microscopy. In addition to endogenous SC35 immunostaining, SC35-YFP was ectopically expressed by transient transfections. Scale bar: 5 µm.

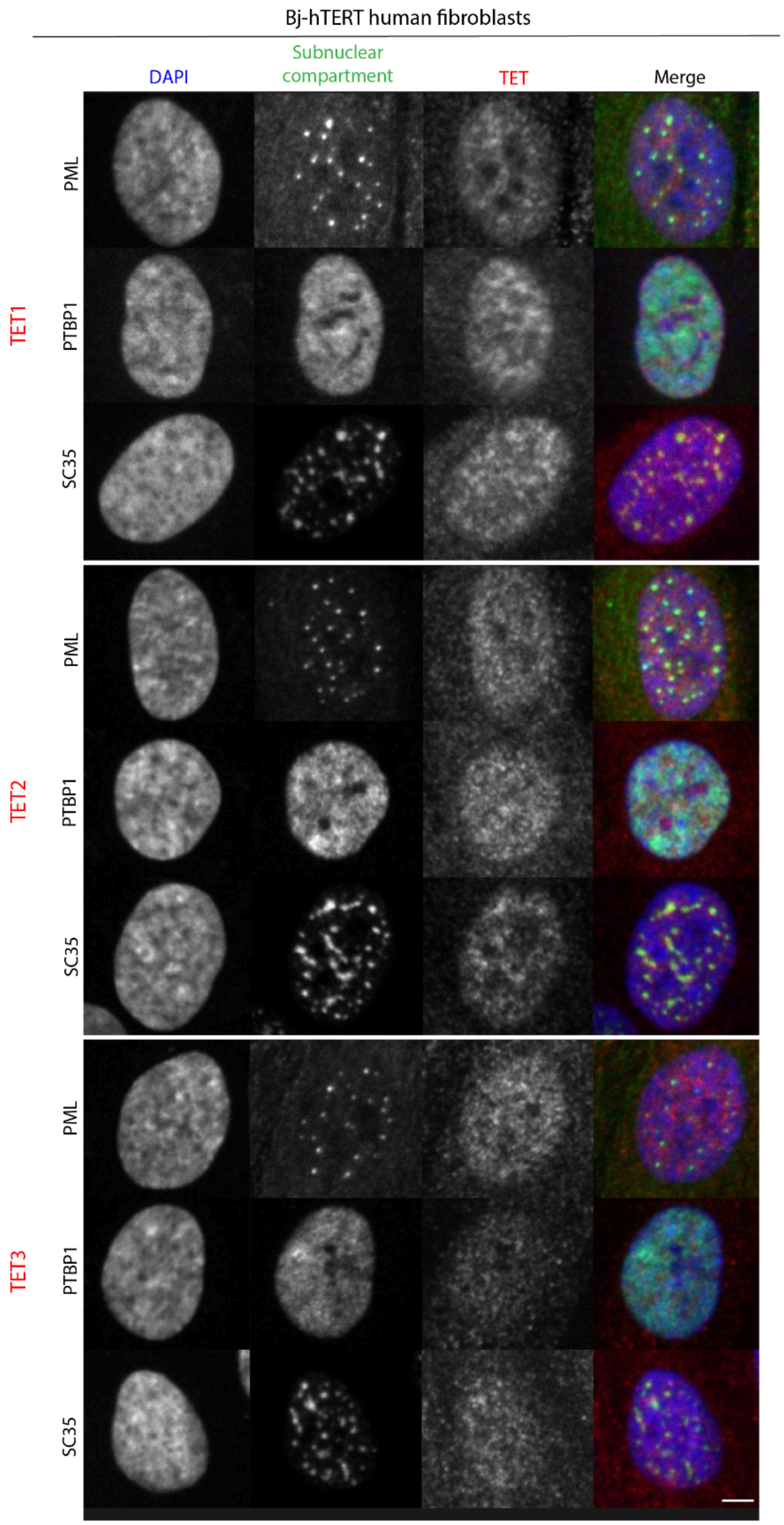

**Figure S4. TET1, TET2, and TET3 localization at splicing speckles visualized with SC35 in human cells.** Bj-hTERT cells (human foreskin fibroblasts immortalized through the ectopic expression of human telomerase reverse transcriptase (hTERT)) were immunostained for endogenous TET proteins and SC35 and imaged by confocal microscopy. In addition to endogenous SC35 immunostaining, PTBP1 (Polypyrimidine Tract Binding Protein 1) involved in alternative splicing regulation, and PML (Promyelocytic Leukemia Protein) involved in nuclear body formation, were stained. Scale bar: 5 μm.

Murine testicular cells

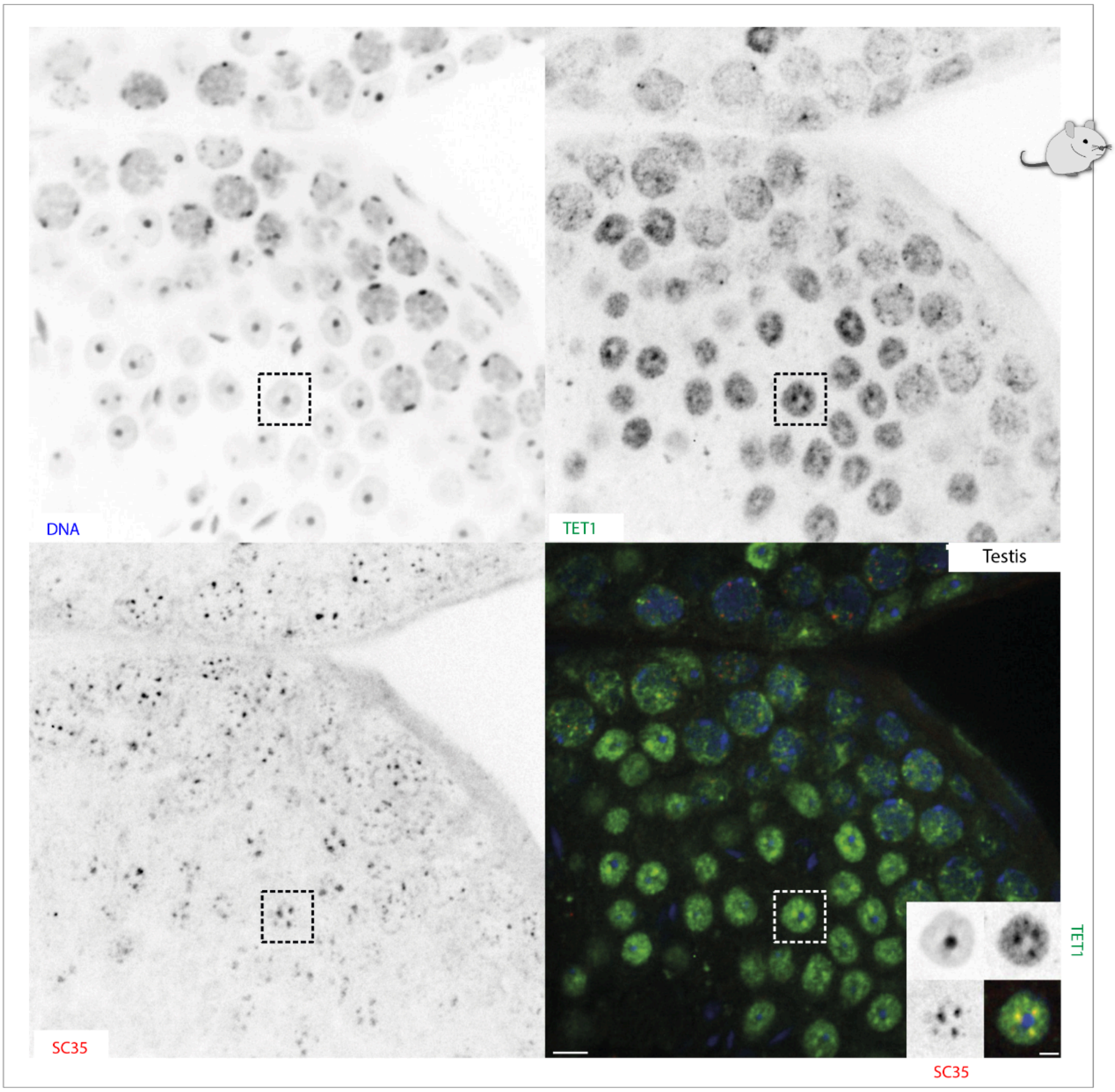

**Figure S5. TET1 localizes to SC35-positive speckles in murine testicular cells.** Paraffin-sections from BALB/c testes were immunostained with antibodies against TET1 and SC35. DNA was counterstained with DAPI. The blowout on the bottom right shows a selected cell and the subnuclear localization of TET1 and SC35 proteins. Stainings for other tissues are shown in Fig. S6 and S7. Scale bar: 5 µm.

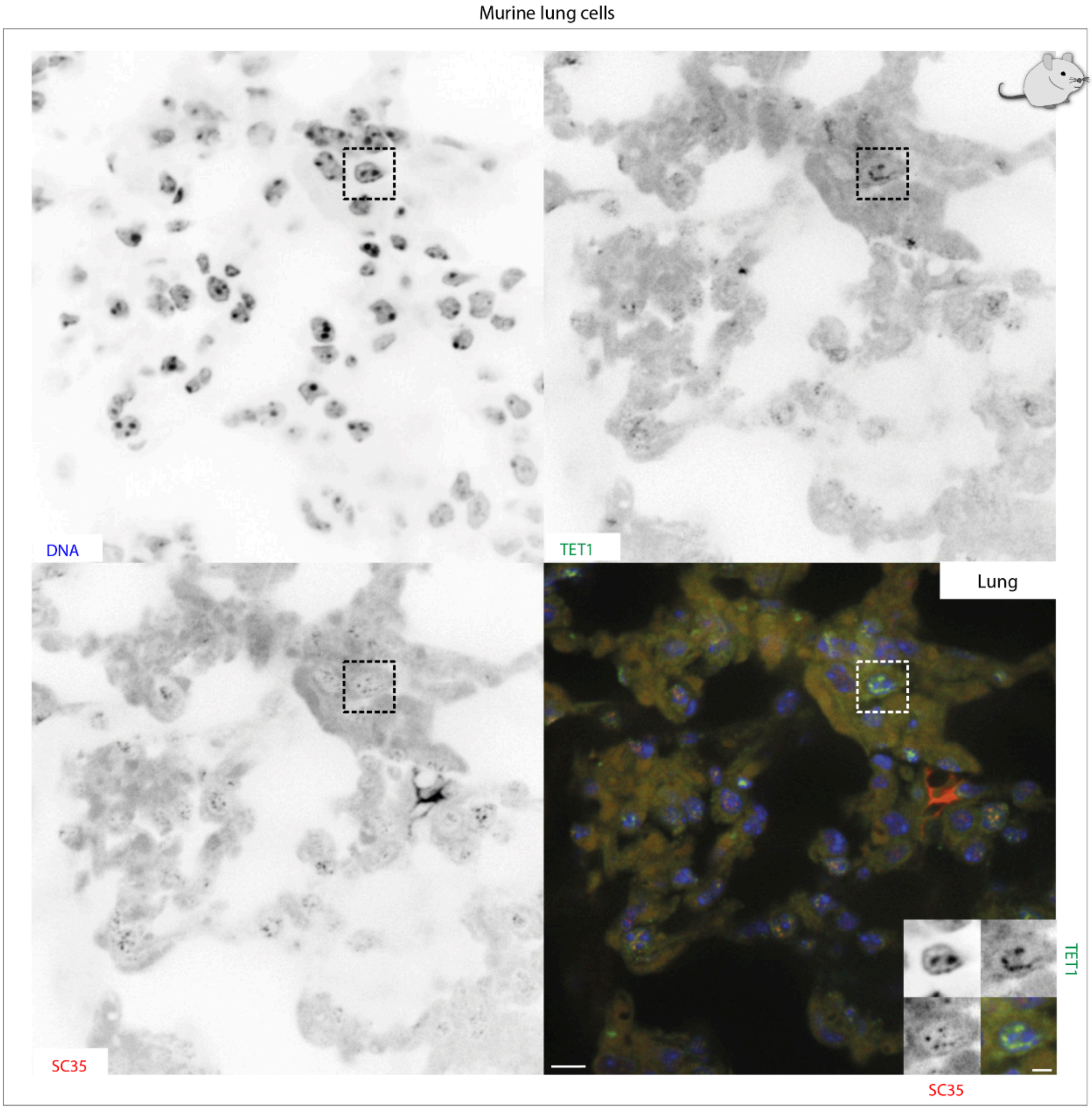

**Figure S6. TET1 localizes to SC35-positive speckles in murine lung cells.** Paraffin sections were immunostained with antibodies against TET1 and SC35. DNA was counterstained with DAPI. The blowout on the bottom right shows a selected cell and the subnuclear localization of TET1 and SC35 proteins. Scale bar: 5 µm.

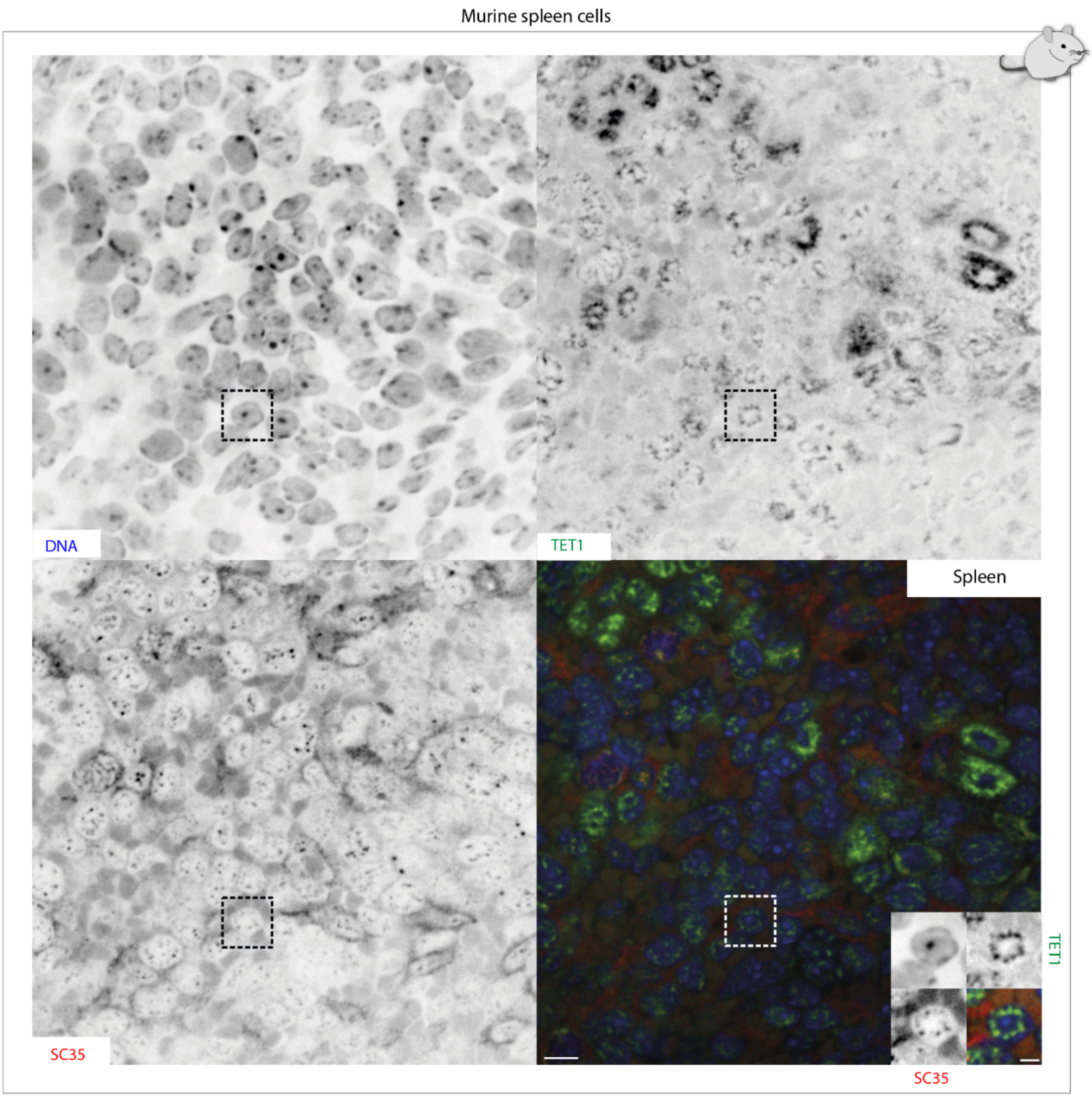

**Figure S7. TET1 localizes to SC35-positive speckles in murine spleen cells.** Paraffin sections were immunostained with antibodies against TET1 and SC35. DNA was counterstained with DAPI. The blowout on the bottom right shows a selected cell and the subnuclear localization of TET1 and SC35 proteins. Scale bar: 5 µm.

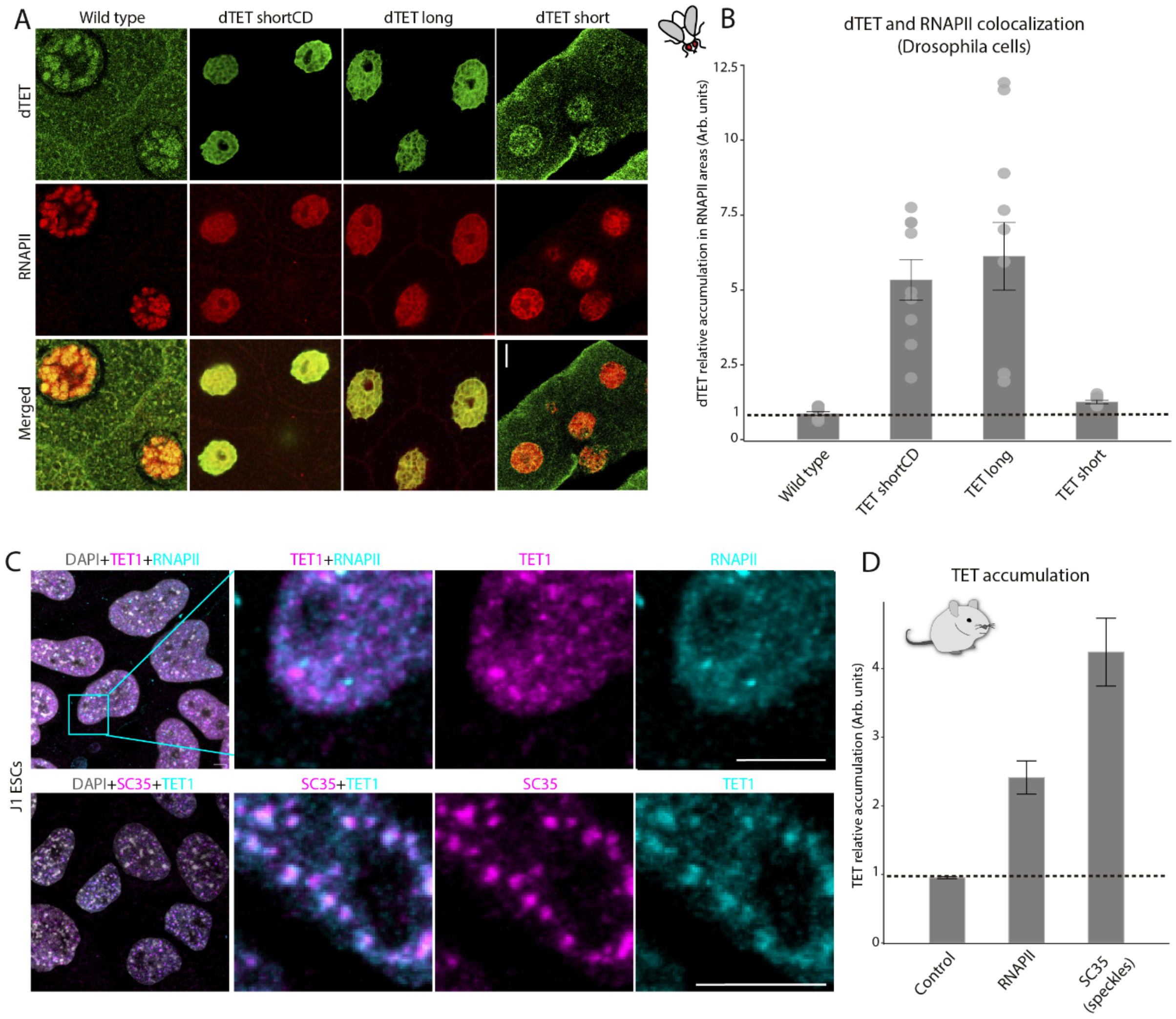

**Figure S8. TET1 colocalizes with SC35 and RNAPII in Drosophila and mouse cells. (A)** HA-tagged short, long, and short catalytically dead mutant (shortCD) TET isoforms were expressed from UAS inserts by elavC155GAL4 in Drosophila cells. Drosophila third instar larvae salivary glands were stained with anti-HA antibodies to visualize TET-tagged proteins and elongating RNAPII (phosphoSer2) antibodies. Scale bar: 15 µm. **(B)** Colocalization of TET1 and RNAPII was quantified by measuring TET relative accumulation in RNAPII areas as performed in Fig. 3C. Barplot shows the result of this analysis, where the dashed black line separates values above 1, indicating colocalization. **(C)** Representative confocal images of J1 mESCs immunostaining for TET1 and RNAPII or SC35. The magnified areas show both the TET1 and RNAPII/SC35 signal and their overlap. Scale bar: 5 µm. **(D)** Colocalization was quantified by image analysis as described in (B) and is shown as a barplot, with the dashed black line separating values above 1.

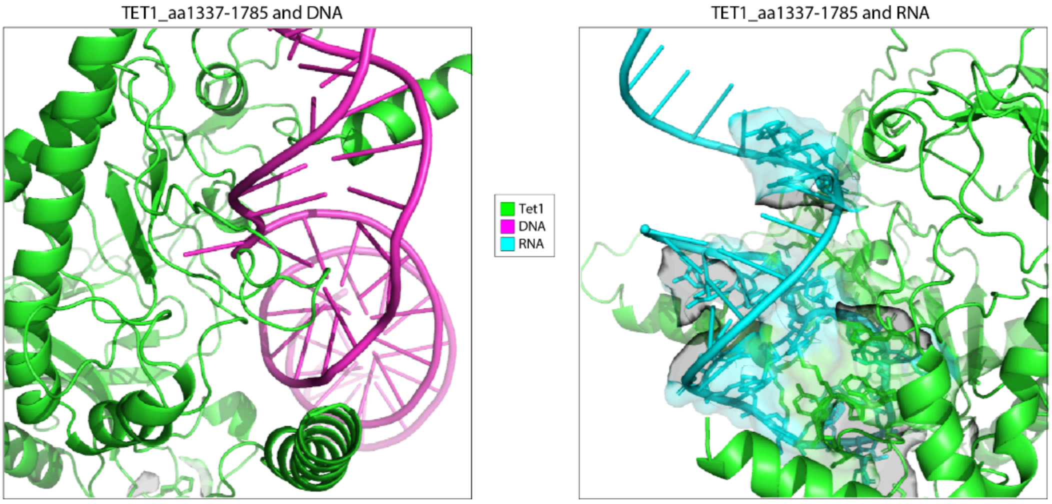

**Figure S9. Structural modeling of TET1-CD interaction with DNA and RNA.** Magnification of the structural models generated with AlphaFold 3 for TET1-CD and DNA interaction (left) versus RNA (right). As a DNA sequence, a region of the 5' UTR LINE1 promoter (5'-GCGCACCTTCCCTGTAAGAGAGCTTGCCAGCAGAGAGTGCTCTGA-3') was chosen based on previous studies. As an RNA sequence, a short region located in an intron/exon limit was selected (5'-AAAACAUAAAGAAAGGCGUGAGUCUAUGGGA-3'). The sequence used was obtained from the splicing reporters used in this study (spliceable luciferase).

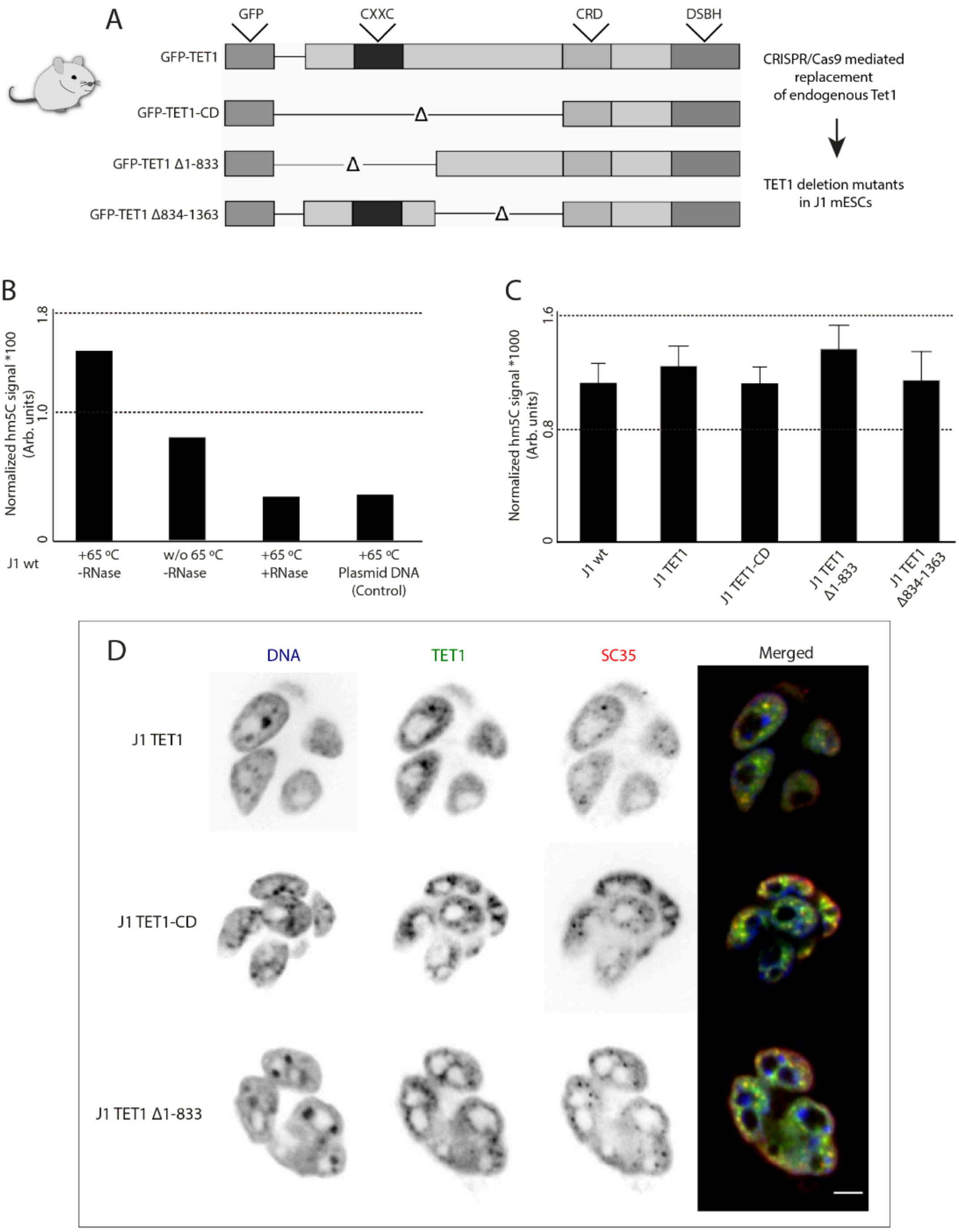

**Figure S10. The catalytic domain of TET1 oxidates m5C to hm5C in RNA secondary structures and localizes in splicing speckles. (A)** Scheme showing GFP-tagged TET1 deletion mutant cDNAs used for the genetic edition of J1 wild-type mESCs by CRISPR/Cas9 technology. The full coding sequence of the mutants without introns replaced the TET1 genomic sequence. The cell lines are shown in Supplementary Table S2. **(B)** Total RNA slot blot analysis from J1 wild-type ESCs and image analysis quantification. Barplots showing the quantification for different samples and treatments: RNA boiled at 65°C without RNase A treatment, RNA sample not boiled and not treated with RNase A, RNA sample boiled at 65°C and treated with RNase A, and plasmid DNA digested with EcoRI and HindIII boiled at 65°C as control. The y-axis shows the hm5C signal in arbitrary units normalized to the methylene blue signal. **(C)** Quantification of slot blot analysis of total RNA isolated from J1 wild-type ESCs and ESCs expressing different TET1 deletion mutants. The y-axis shows the averaged hm5C signal in arbitrary units normalized to the RNA concentration determined by methylene blue intensity. Error bars represent the standard deviation. Experiments were performed as biological triplicates. **(D)** Representative images of immunofluorescence staining for TET1 and SC35 in the different J1 mESCs TET1 deletion mutants. Scale bar: 10 µm.

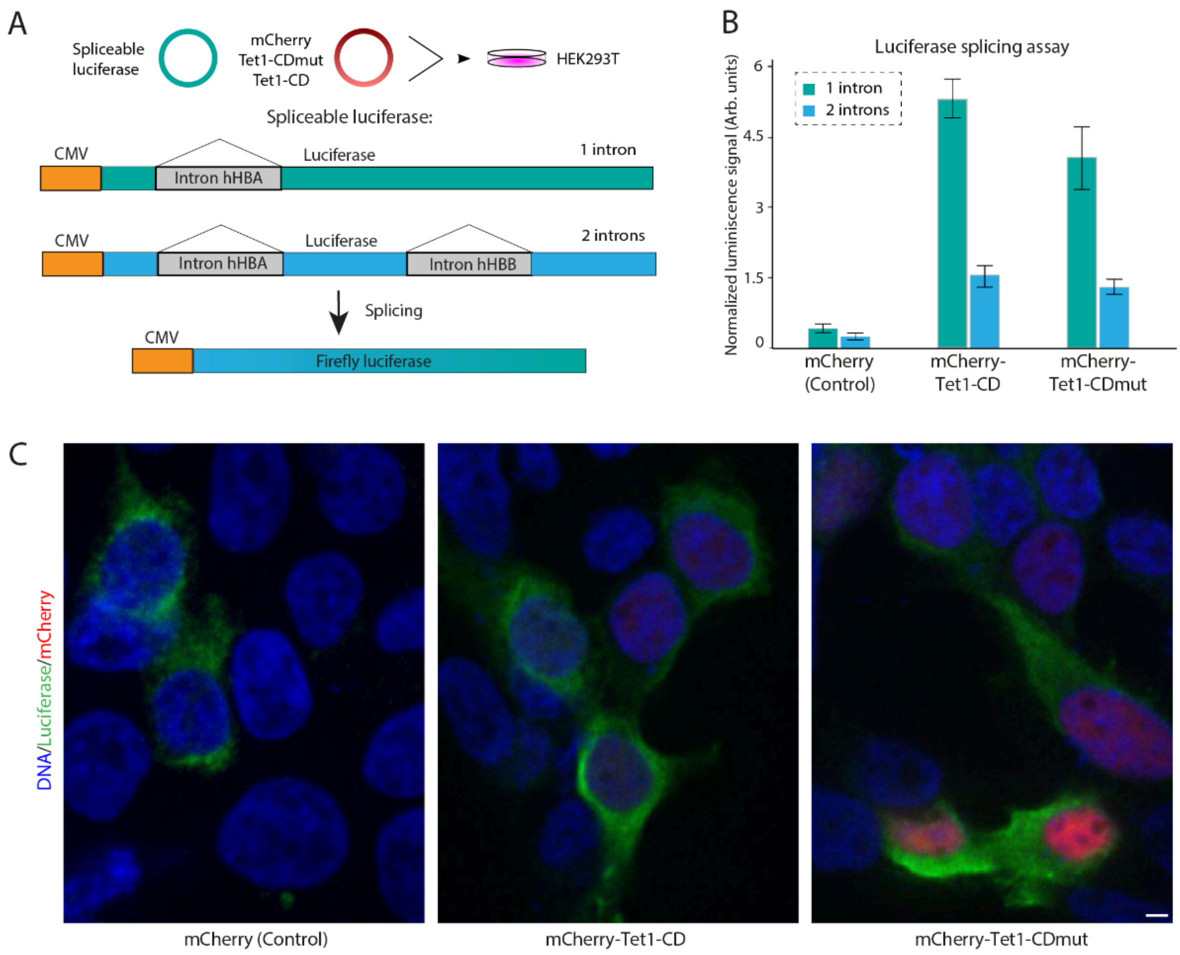

**Figure S11. TET1 promotes splicing independent of its catalytic activity. (A)** A splicing reporter assay with spliceable firefly luciferase was used to study the role of TET1-CD and TET1-CDm (catalytic dead mutant) in alternative splicing. The diagram shows the pipeline of the experiment and the structure of the luciferase constructs used: spliceable luciferase with two introns (human hemoglobin alpha [hHBA] and human hemoglobin beta [hHBB]) is expressed under the control of a CMV promoter. Alternatively, spliceable luciferase with one intron (hHBA) is also used. **(B)** Barplot showing quantification of the splicing reporter assay by measuring luminescence signal. The results for each construct co-transfected with luciferase genes either containing one (green) or two introns (blue) are shown, and mCherry was used as a control. The quantified luminescence signal in arbitrary units was normalized to the fluorescent signal of mCherry for each condition. Error bars represent the standard deviation. Experiments were performed as biological triplicates. Representative images are shown in **(C)**. Scale bar: 5 µm.

### Supplementary tables

#### Supplementary Table S1: Plasmids

| Name | pc number* | Fluorophore/tag | Gene species | Promoter | Reference |
|---|---|---|---|---|---|
| pEGFP-N1 | 0713 | GFP | *Aequorea victoria* | CMV | Clontech |
| pGFP-mTet1 | 2271 | GFP | *Mus musculus* | CAG | (Frauer et al., 2011) |
| pGFP-mTet2 | 2272 | GFP | *Mus musculus* | CAG | (Bauer et al., 2015) |
| pGFP-mTet3 | 2273 | GFP | *Mus musculus* | CAG | (Liu et al., 2013) |
| pCherry-C1 | 2387 | mCherry | *Discosoma sp.* | CMV | (Becker et al., 2016) |
| pCherry-mTet1-CD | 2547 | mCherry | *Mus musculus* | CAG | (Ludwig et al., 2017) |
| pCherry-mTet1-CDmut | 2815 | mCherry | *Mus musculus* | CAG | (Ludwig et al., 2017) |
| pCherry-mTet2-CD | 3338 | mCherry | *Mus musculus* | CAG | (Zhang et al., 2017b, 2017c) |
| pCherry-mTet3-CD | 3339 | mCherry | *Mus musculus* | CAG | (Ludwig et al., 2017) |
| pCAG-GFP-Tet1CD | 2315 | GFP | *Mus musculus* | CAG | (Spruijt et al., 2013) |
| pCAG-GFP-Tet2CD | 2316 | GFP | *Mus musculus* | CAG | (Arroyo et al., 2022; Zhang et al., 2017b, 2017c) |
| pCAG-GFP-Tet3CD | 2309 | GFP | *Mus musculus* | CAG | (Arroyo et al., 2022; Ludwig et al., 2017) |
| pU2AF35-FLAG | 3967 | Flag | *Mus musculus* | CMV | Gift from Florian Heyd (Gama-Carvalho et al., 1997; Herdt et al., 2020) |
| pU2AF65-HA | 3968 | HA | *Mus musculus* | CMV | Gift from Florian Heyd (Gama-Carvalho et al., 1997; Herdt et al., 2020) |
| pEYFP-SC35 | 1202 | YFP | *Mus musculus* | CMV | (Richter et al., 2005) |
| pmCherry-SC35 | 3332 | mCherry | *Mus musculus* | CMV | This study |
| pMCP-EGFP-LacI | 4555 | GFP | Artificial sequence | CMV | (Duan et al., 2021) |
| pms2-PABCP1-mCherry | 4573 | mCherry | *Mus musculus* | CMV | (Duan et al., 2021) |
| pPABPC1-mCherry | 4588 | mCherry | *Mus musculus* | CMV | (Duan et al., 2021) |
| pUC18-MINX-M3 | 3902 | - | Artificial sequence | T7 | Gift from Lührmann Lab (Deckert et al., 2006; Zhang et al., 2017a; Zillmann et al., 1988) |
| hnRNP-DL-EGFP_ECFP - Splicing reporter NMD | 3351 | GFP | Synthetic construct | CMV | This study |
| pWHE200-bg - Splicing reporter luciferase | 3335 | luciferase | Synthetic construct | CMV | Generated by C. Berens lab, provided by Suess Lab (Vogel et al., 2018) |
| pWHE237-bg - Splicing reporter luciferase | 3336 | luciferase | Synthetic construct | CMV | Generated by C. Berens lab, provided by Suess Lab (Vogel et al., 2018) |
| pWHE237mod-WT - Splicing reporter luciferase | 3337 | luciferase | Synthetic construct | CMV | Suess Lab (Vogel et al., 2018) |

*pc number: plasmid collection number

#### Supplementary Table S2: Mammalian cell lines

| Name | Species | Type | Genotype | Gender | Reference/Source |
|---|---|---|---|---|---|
| HEK293-EBNA | *Homo sapiens* | Embryonic kidney | wildtype | female | CVCL_6974 Invitrogen; Paisley, UK |
| ES v6.5 wt | *Mus musculus* | Embryonic stem cells | wildtype | male | (Dawlaty et al., 2014) |
| ES v6.5 Tet TKO | *Mus musculus* | Embryonic stem cells | Tet triple knockout | | (Dawlaty et al., 2014; Zhang et al., 2017b) |
| ES J1 wt | *Mus musculus* | Embryonic stem cells | wildtype | male | (Li et al., 1992) |
| ES J1 Dnmt 1/3a/3b TKO | *Mus musculus* | Embryonic stem cells | Dnmt triple knockout | male | (Okano et al., 1999; Tsumura et al., 2006) |
| ES J1 Tet1 Δ 834-1363 | *Mus musculus* | Embryonic stem cells | GFP–Tet1Δ834-1363 (endogenous Tet1 deletion) | male | (Mulholland et al., 2015) |
| ES J1 Tet1 Δ1- 833 | *Mus musculus* | Embryonic stem cells | GFP- Tet1Δ1- 833 | male | (Mulholland et al., 2015) |

| | | | | | |
|---|---|---|---|---|---|
| ES J1 Tet1 CD | *Mus musculus* | Embryonic stem cells | GFP-Tet1CD (endogenous Tet1 deletion) | male | (Mulholland et al., 2015) |
| ES J1 Tet1 cDNA F4 | *Mus musculus* | Embryonic stem cells | GFP-Tet1 full-length (endogenous Tet1 deletion) | male | (Mulholland et al., 2015) |
| MEF W8 | *Mus musculus* | Mouse embryonic fibroblasts | wildtype | male | (Peters et al., 2001) (Jenuwein Lab, Freiburg, Germany) |
| MTF wt line 3 | *Mus musculus* | Mouse tail fibroblasts | wildtype | female | (Guy et al., 2001) (Bird Lab, Edinburgh, UK) |
| BHK clone 2 | Mesocricet us auratus | Baby hamster kidney fibroblast | Stable lac-operator array | male | (Tsukamoto et al., 2000) |
| Bj-hTERT | *Homo sapiens* | Skin fibroblast | wildtype | male | Gift from Mathias Rosenfeldt (Aladjem and Fanning, 2004; Bodnar et al., 1998) |
| HeLa | *Homo sapiens* | Cervix epithelium carcinoma | wildtype | female | Stuart Orkin´s lab (HMS) |

##### Supplementary Table S3: Primary and secondary antibodies

| Reactivity | Host* | Dilution | Application** | Cat.No. / Clone / ID | Provider / Reference |
|---|---|---|---|---|---|
| α-TET1 | Rat (mAb) | 1:10 / 1:2 | IF | Clone 5D8 Clone 4H7 | Hybridoma supernatant (Bauer et al., 2015) |
| α-TET2 | Rat (mAb) | 1:10 / 1:2 | IF | Clone 9F7 | Hybridoma supernatant (Bauer et al., 2015) |
| α-TET3 | Rat (mAb) | 1:10 / 1:2 | IF | Clone 11B6 | Hybridoma supernatant (Bauer et al., 2015) |
| α-GFP | Rat (mAb) | 1:1000 | WB | Clone 3H9 | Chromotek, Planegg-Martinsried, Germany |
| GFP binder | nanobody | 1 mg/mL | coIP/mass spectrometry | - | (Rothbauer et al., 2008) |
| α-RFP | Rat (mAb) | 1:200 | WB | Clone 5F8 | (Rottach et al., 2008) |
| α-hemagglutinin (HA) | Mouse (mAb) | 1:200 | WB/IF | Clone 12CA5 | Hybridoma supernatant (Wilson et al., 1984) |
| α-hemagglutinin (HA) | Rabbit | 1:1000 | IF Drosophila cells | | Sigma-Aldrich Chemie |
| α-FLAG Tag | Mouse (mAb) | 1:200 | WB/IF | M2 SLBJ7864V | Sigma-Aldrich Chemie |
| α-Splicing Factor SC-35 | Mouse (mAb) | 1:1000 | IF | clone S-4045 | Sigma-Aldrich Chemie |
| α-Oct4 | rabbit | 1:100 | IF | ab19857 | Abcam, Cambridge, UK |
| α-RNA polymerase II CTD repeat YSPTSPS (phosphoSer5) | Mouse (mAb) | 1:1000 | IF | ab 5408 | Invitrogen |
| α-RNA Pol II (phosphoSer2) | Rat (pAb) | 1:1000 | IF Drosophila cells | clone 3E10, 04-1571 | Merck-Millipore |
| α-5hmC | Rabbit (pAb) | 1:250 | Slot blot | 39769 | Active Motif, La Hulpe, Belgium |
| α-PML (PG-M3) | Mouse (mAb) | 1:100 | IF | J1904 | Santa Cruz Biotechnology |
| α-PTBP1 | Rabbit (mAb) | 1:100 | IF | ab133734 | Abcam, Cambridge, UK |
| α-Mouse IgG Alexa Fluor®555 | Donkey (pAb) | 1:250 | IF | A31570 | Invitrogen, California, USA |
| α-Mouse IgG Cy3 | Donkey (pAb) | 1:250 | IF | 715-166-151 | Jackson Immuno Research, Pennsylvania, USA |
| α-Rat IgG DyLight®488 | Donkey (pAb) | 1:250 | IF | 712-546-153 | Jackson Immuno Research, Pennsylvania, USA |
| α-Rat IgG Alexa Fluor ®488 | Donkey (pAb) | 1:250 | IF | 712-545-153 | Jackson Immuno Research, Pennsylvania, USA |
| α-Rat IgG Cy5 | Donkey (pAb) | 1:250 | IF | 712-175-153 | Jackson Immuno Research, Pennsylvania, USA |
| α-Rat IgG Cy3 | Donkey | 1:250 | IF | 711-165-152 | Jackson Immuno Research, |

| | | | | | |
|---|---|---|---|---|---|
| | (pAb) | | | | Pennsylvania, USA |
| α-rabbit IgG (H+L) Alexa Fluor 488-conjugated | Goat (pAb) | 1:500 | IF | A-11008 | Thermo Fisher Scientific, Waltham, MA, USA |
| α-mouse IgG (H+L) Alexa Fluor 488-conjugated | Goat (pAb) | 1:500 | IF | A-11001 | Thermo Fisher Scientific, Waltham, MA, USA |
| α-rabbit IgG (H+L) Alexa Fluor 594-conjugated | Goat (pAb) | 1:250 | IF | R37117 | Thermo Fisher Scientific, Waltham, MA, USA |
| α-Rat IgG Alexa Fluor 647-conjugated | Goat (pAb) | 1:250 | IF | A-21247 | Thermo Fisher Scientific, Waltham, MA, USA |
| α-Rat IgG (H+L) HRP***-conjugated | Goat (pAb) | 1:5000 | WB | A9037 | Sigma-Aldrich, St Louis, MO, USA |
| α-Mouse IgG (H+L) HRP***-conjugated | Sheep (pAb) | 1:5000 | WB | NA931 | GE Healthcare, Chicago, Il, USA |

*mAb: monoclonal; pAb: polyclonal; **IF: immunofluorescence, WB: western blot; ***HRP: horseradish peroxidase

##### Supplementary Table S4: Imaging systems

| Device | Light Sources | Filters (ex & em [nm])* | Objectives/ Lenses | Detection system | Application |
|---|---|---|---|---|---|
| Ultra-View VoX spinning disc on an Inverted Nikon Ti-E microscope; PerkinElmer Life Sciences, UK | Solid state Diode lasers (405 nm, 488 nm, 561 nm, 640 nm) | 405/488/568/640 **<br>405: 415–475<br>488: 505–549<br>561: 580–650<br>640: 664–754 | Oil immersion 60x Plan-Apochromat (NA 1.49)<br>Oil immersion 100x Plan-Apochromat (NA 1.49) | cooled 14-bit Hamamatsu® C9100-50 EMCCD | time-lapse microscopy & confocal Z-stack imaging |
| Leica SP5 II Confocal point scanner | 405 nm Diode Laser 50 mW<br>488 nm Argon ion laser<br>458 nm ~5mW<br>476 nm ~5mW<br>488 nm ~20mW<br>496 nm ~5mW<br>514 nm ~20mW<br>561 nm DPSS 50 mW<br>633nm HeNe gas laser 20 mW | DAPI:<br>ex. 420/30 em. 465/20<br><br>FITC:<br>ex. 495/15 em. 530/30<br><br>Rhod:<br>ex. 570/20 em. 640/40<br><br>FITC/Rhod combi filter | HC PL APO 10x / 0.4 CS<br><br>HCX PL APO 40x / 1.3 oil CS<br><br>HCX PL APO 63x / 1.4-0.6 oil lambda blue<br><br>HCX PL APO 100x / 1.44 oil Corr CS | spectral from 400-800 selectivity 0.6-2 nm<br>galvano scanner: up to 1400 Hz<br>resonance scanner: up to 8000 Hz, up to 250 frames per second with 512x512 | Time-lapse microscopy & confocal Z-stack imaging |
| Operetta high content screening microscope; PerkinElmer Life Sciences, UK | Xenon fiber optic Light source, 300 W, 360 – 640 nm continuous spectrum | 405: 360-400 & 410–480<br>488: 460-490 & 500–550<br>561: 560-580 & 590–640 | 20x or 40x air (0.45 NA and 0.95 NA) long WD*** | 14-bit Jenoptik CMOS | high content screening microscopy |
| Amersham AI600 Imager; GE Healthcare, Chicago, Il, USA | UV transillumination light: 312 nm | EtBr: 312 & 585-625 | Large aperture f/0.85 FUJINON™ | 16-bit Peltier cooled Fujifilm Super CCD | HRP stained blot imaging |

*ex: excitation & em: emission, ** dichroic specification, *** WD: working distance

##### Supplementary Table S5: Statistics

| Figure | Sample | n/replicates | Mean | StDev | p-value |
|---|---|---|---|---|---|
| **1A-C** | See Supplementary Data 1 | | | | |
| **2C** | See Supplementary Data 2 | | | | |
| **3A** | J1 (TET&DAPI) | 10/2 | 1.221875845 | 0.18423886 | NA |
| | J1 (TET&SC35) | 10/2 | 2.419255459 | 0.18133659 | |
| | EB (TET&DAPI) | 10/2 | 0.845532755 | 0.23053236 | |
| | EB (TET&SC35) | 10/2 | 1.989337709 | 0.246316896 | |
| | MEF (TET&DAPI) | 10/2 | 0.675793258 | 0.137883411 | |
| | MEF (TET&SC35) | 10/2 | 1.543048949 | 0.076502812 | |
| | MTF (TET&DAPI) | 10/2 | 0.720262662 | 0.154411838 | |
| | MTF (TET&SC35) | 10/2 | 1.786633442 | 0.147527388 | |
| **3B** | MTF (TET&DAPI) | 10/2 | 0.432843 | 0.1934187 | NA |

| | | | | | |
|---|---|---|---|---|---|
| | MTF (TET&U2AF1)<br>MTF (TET&DAPI)<br>MTF (TET&U2AF1) | 10/2<br>10/2<br>10/2 | 1.769327<br>0.419264<br>1.891072 | 0.2681941<br>0.1876103<br>0.2902541 | |
| **3D** | dTET short<br>dTET long<br>dTET shortCD<br>Wild-type (CantonS) | 5/2<br>5/2<br>5/2<br>5/2 | 1.608761001<br>1.607218392<br>1.489584822<br>1.039347709 | 0.198854875<br>0.033577065<br>0.190533916<br>0.043284972 | NA |
| **4A-D** | See Supplementary Data S3-4 | | | | |
| **5A** | Untreated<br>-Triton+RNase<br>+Triton-RNase<br>+Triton+RNase | 13566/3<br>10060/3<br>11799/3<br>6203/3 | 1<br>0.72934<br>0.92823<br>0.26342 | 0<br>0.14823<br>0.00173<br>0.17329 | NA |
| **5C** | (1) Negative control<br>(2) Positive control<br>(3) RNA Trap + mRNA mimic | 24/2<br>23/2<br>37/2 | 1.059673486<br>3.422852525<br>2.447521656 | 0.1633333<br>1.471513136<br>0.899997807 | (1-2) 8.417e-12<br>(1-3) 2.2e-16<br>(2-3) 0.004573 |
| **5E** | See Supplementary Data 2 | | | | |
| **6B** | TET Triple KO<br>WT+Cherry<br>TKO+Cherry<br>TKO+TET1CDm<br>TKO+TET1CD<br>TKO+TET2CD<br>TKO+TET3CD | <br>13362/2<br>3558/2<br>21989/2<br>23120/2<br>14513/2<br>21153/2 | <br>1.38769884<br>1.071902182<br>5.68629579<br>4.782332972<br>4.657309325<br>4.654218624 | <br>0.904570142<br>0.63286406<br>2.908236827<br>2.616347675<br>2.958514821<br>2.701217227 | NA |
| **6C** | Dnmt Triple KO<br>WT+Cherry<br>TKO+Cherry<br>TKO+TET1CDm<br>TKO+TET1CD<br>TKO+TET2CD<br>TKO+TET3CD | <br>8933/2<br>29939/2<br>11860/2<br>15505/2<br>2852/2<br>14303/2 | <br>3.436116618<br>2.560897662<br>4.807776619<br>4.744117214<br>4.173808474<br>4.870746262 | <br>2.362970511<br>1.769923453<br>2.819110378<br>2.660777855<br>2.816790199<br>2.652516561 | NA |
| **S8B** | Wild type<br>TET shortCD<br>TET long<br>TET short | 10/2<br>9/2<br>11/2<br>8/2 | 0.869797722<br>5.335722866<br>6.123324018<br>1.257700419 | 0.208917705<br>2.04129062<br>3.757580461<br>0.162348096 | NA |
| **S8D** | Control<br>RNAPII<br>SC35 (speckles) | 12/2<br>14/2<br>23/2 | 0.954414887<br>2.41385661<br>4.242922308 | 0.047514536<br>0.873790948<br>2.208899833 | NA |
| **S10B** | +65 ºC -RNase<br>w/o 65 ºC -RNase<br>+65 ºC +RNase<br>+65 ºC Plasmid DNA (control) | /1<br>/1<br>/1<br>/1 | 1.5396<br>0.8293<br>0.2945<br>0.3021 | -<br>-<br>-<br>- | NA |
| **S10C** | J1 wt<br>J1 TET1<br>J1 TET1-CD<br>J1 TET1-Δ1-833<br>J1 TET1-Δ8341363 | /3<br>/3<br>/3<br>/3<br>/3 | 1.1256<br>1.3573<br>1.1102<br>1.3814<br>1.0991 | 0.2156<br>0.2573<br>0.1985<br>0.3012<br>0.3928 | NA |
| **S11B** | mCherry (1 intron)<br>mCherry (2 introns)<br>mCherry-TET1-CD (1 intron)<br>mCherry-TET1-CD (2 introns)<br>mCherry-TET1-CDmut (1 intron)<br>mCherry-TET1-CDmut (2 introns) | >190377/3<br>>89627/3<br>>168140/3<br>>70563/3<br>>81505/3<br>>161862/3 | 0.2301<br>0.1683<br>5.2638<br>1.6112<br>3.6183<br>1.2196 | 0.1134<br>0.1021<br>0.7632<br>0.3198<br>0.9856<br>0.2659 | NA |
